# Adherens junction serves to generate cryptic lamellipodia required for collective migration of epithelial cells

**DOI:** 10.1101/2020.04.20.052118

**Authors:** Masayuki Ozawa, Sylvain Hiver, Takaki Yamamoto, Tatsuo Shibata, Srigokul Upadhyayula, Yuko Mimori-Kiyosue, Masatoshi Takeichi

## Abstract

Collective migration of epithelial cells plays crucial roles in various biological processes such as cancer invasion. In migrating epithelial sheets, leader cells form lamellipodia to advance, and follower cells also form similar motile apparatus at cell-cell boundaries, which are called cryptic lamellipodia (c-lamellipodia). Using adenocarcinoma-derived epithelial cells, we investigated how c-lamellipodia are generated, and found that they sporadically grew from Ecadherin-based adherens junctions (AJs). WAVE and Arp2/3 complexes were localized along the AJs, and silencing them not only interfered with c-lamellipodia formation but also prevented follower cells from trailing the leaders. Disruption of AJs by removing αE-catenin resulted in uncontrolled c-lamellipodia growth, and this was brought about by myosin II activation and the resultant contraction of AJ-associated actomyosin cables. Additional observations indicated that c-lamellipodia tended to grow at mechanically weak sites of the junction. We conclude that AJs not only tie cells together but also generate c-lamellipodia by recruiting actin regulators, enabling epithelial cells to undergo ordered collective migration.

## Introduction

Animal cells migrate as a collective in many morphogenetic processes, as well as in pathological events such as cancer invasion (Cheung and Ewald, 2014; De Pascalis and Etienne-Manneville, 2017; Friedl and Alexander, 2011; Friedl and Gilmour, 2009). It is therefore important to understand why cells move together rather than as single cells, and how the movement of individual cells is controlled and coordinated to allow their collective migration. Various types of cells require cadherin-mediated cell-cell adhesion for their orderly migration not only *in vivo* (Cai et al., 2014b; Niewiadomska et al., 1999), but also *in vitro* (Camand et al., 2012; Desai et al., 2009; Dupin et al., 2009; Ladoux and Mege, 2017; Mayor and Etienne-Manneville, 2016). This suggests that cadherins regulate cell behavior that is necessary for collective migration. However, the precise mechanisms of how epithelial cells require cadherins for their collective migration are not yet known.

Cells of ‘simple epithelia’ are connected to each other via a ‘junctional complex,’ which consists of a tight junction (TJ), adherens junction (AJ, formally zonula adherens) and desmosome, at the apical-most end of cell-cell contacts (Farquhar and Palade, 1963). Due to the observation that the TJ and AJ are closely adjoined to one another, this set of junctions is often called the apical junctional complex (AJC) (Anderson et al., 2004; Vogelmann and Nelson, 2005). The AJC associates with a bundle of actin cables, called the circumferential actin belt or cable, which encircles individual cells at their apical ends, resulting in a honeycomb-like pattern of distribution. Below the junctional complex, non-specialized junctions, conveniently termed the lateral cell-cell contacts (LCs), extend to the basal end of the cell, which actually occupies most areas of the cell junction. E-cadherin is a main adhesion receptor at the AJ of epithelial cells, which also functions at LCs. It binds β-catenin or plakoglobin and in turn αE-catenin, forming the cadherin-catenin complex (CCC). αEcatenin directly interacts with F-actin, or indirectly does so via binding to vinculin. In the absence of αE-catenin, E-cadherin is unable to maintain the AJC, indicating that the interaction of CCC with F-actin is crucial for the epithelial-specific junction organization (Mege and Ishiyama, 2017; Takeichi, 2014).

The lamellipodium is a major structure in cell motility. At its front edge, actin polymerization is initiated, and the resultant actin filaments grow backward forming meshwork under the control of numerous regulators, including Rac1 and its effectors (Ridley, 2015). This actin regulation generates a force for the cellular margin to advance, accompanied by dynamic membrane ruffling. When cells migrate as a collective, leader cells, which occupy the front edge of a cell sheet, generate lamellipodia to move forward, being trailed by follower cells (Haeger et al., 2015; Omelchenko et al., 2003). The followers also organize protrusions or lamellipodium-like structures, called cryptic lamellipodium (c-lamellipodium), most likely in order to chase the leaders (Farooqui and Fenteany, 2005). Similar structures related to cell movement are also detectable when epithelial cells move without any leader cells (Barlan et al., 2017; Krndija et al., 2019; Squarr et al., 2016).

Cadherin-mediated cell-cell contacts are known as a mechanism to interfere with cell motility, particularly in the process of contact inhibition of cell locomotion (Roycroft and Mayor, 2016; Theveneau et al., 2010). Such reported role of cadherins seemingly contradicts the observation that cryptic lamellipodia still form at cell-cell boundaries. In the present study, we investigated how epithelial cells manage their motility at cell-cell contact zones, using adenocarcinoma-derived cell lines. Our observations indicate that AJs not only function to tie cells together and prevent random movement, but they also serve as a site of clamellipodia formation by recruiting the WAVE regulatory complex and its effectors. Thus, we revealed an unexpected function of AJs: to support the migration of epithelial cells as a sheet.

## Results

### Epithelial cells require cell junctions to migrate

To re-examine the role of AJs in epithelial cell migration, we disrupted them by removing the αE-catenin gene (*CTNNA1*) in three adenocarcinoma lines: DLD1, Caco2 (both derived from colon carcinoma), and MKN74 (derived from gastric carcinoma) using the CRISPR/Cas9 method (Fig. S1A). We isolated *CTNNA1*-deleted clones for each line, and subjected them to a classical wound healing assay using collagen-coated or non-coated substrates. The results showed that αE-catenin removal caused an overall delay in wound healing for any cell line (Figs. 1A and 1B), although their initial speed of migration varied from experiment to experiment, probably due to multiple factors that affect wound healing movement (De Pascalis and Etienne-Manneville, 2017). Time-lapse tracking of DLD1 cells at the front margin of their sheets showed that wild-type cells exhibited directed movement, whereas *CTNNA1*-deleted DLD1 (αEcat KO DLD1) cells moved in a non-straight fashion (Fig. 1A and Video 1), explaining why the latter were delayed in wound healing. αE-catenin removal did not significantly alter the proliferation of these cells. For example, in the case of DLD1 and Caco2 cells, the percentages of mitotic cells in culture were 3.42±1.81 and 4.07±1.64 for wild-type (n=12) and αEcat KO DLD1 (n=10, p=0.2), respectively; and 3.43±0.33 and 3.96±0.17 for wild-type (n=6) and *CTNNA1*-deleted (αEcat KO) Caco2 (n=6, p=0.2), respectively. These results indicate that the epithelial cells used here require the cadherin adhesion system for their efficient migration, as shown for other cell types (Mayor and Etienne-Manneville, 2016).

**Figure 1.**
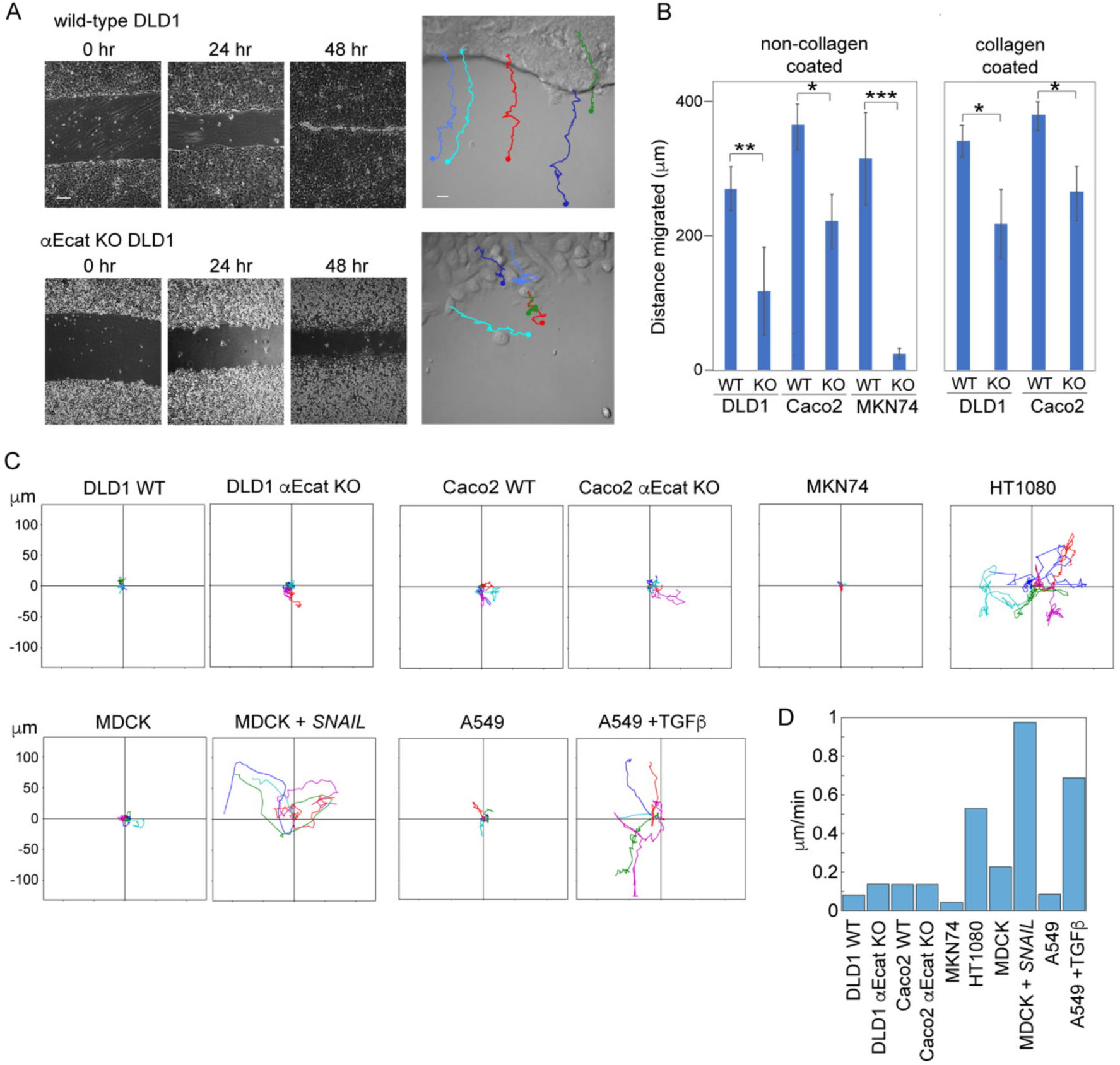
Junction-dependent migration of epithelial cells. (**A**) Wound healing assays of wild-type and *CTNNA1*-deleted (αEcat KO) DLD1 cells. In the right-most panels, trajectories of five independent cells during a 20-h time span are drawn over a montage image of Video 1. Scale bars, 10 μm for the right-most panels and 200 μm for other panels. (**B**) Migration distance of the marginal cells at 24 hrs after scratching of the culture. WT, wild-type cells; KO, αEcat KO cells. Values were obtained at 25 points in a culture photographed with a 4X objective. Three cultures were analyzed. *p<0.05, **p<0.01, ***p<0.001 (t-test). (**C**) Trajectories of singly isolated cells recorded for 20 hrs. MDCK+*SNAIL*, MDCK cells stably transfected with *SNAIL.* A549+TGFb, cells cultured in 5 ng/ml of TGF-β. Values in the vertical and horizontal axes are identical. (**D**) Migration speed of isolated cells. Movies used for C were analyzed to obtain migration speed of each cell by temporal averaging of its instantaneous speed, and then by averaging of the values across the ensemble.

To further study the role of cell-cell adhesion in cell migration, we observed the behavior of singly isolated DLD1 or Caco2 by time-lapse movies. Contrary to the observations using other epithelial types such as keratinocytes (Euteneuer and Schliwa, 1984), isolated DLD1 or Caco2 cells did not show any extensive migration, only movement around a fixed position (Fig. 1C and Video 2). To test whether this is a general property of simple epithelium-derived cells, we examined other cell lines, MKN74, A549 (lung adenocarcinoma) and MDCK cells, and confirmed that they also do not migrate when isolated (Fig. 1C). As mesenchymal cells are known to self-migrate, we examined the effect of epithelial-mesenchymal transition (EMT) on the migration of these cells, using MDCK cells transfected with *SNAIL* cDNA (Ozawa and Kobayashi, 2015), or A549 cells treated with TGF-β (Thiery, 2003). The resultant mesenchyme-like cells became highly migratory, as observed with the fibroblastic line HT1080 (Fig. 1C), exhibiting an increase of their migration speed (Fig. 1D). Thus, these epithelial cells were unable to migrate alone, unless transformed into the mesenchymal type. For further experiments, we chose either of αEcat KO DLD1 or αEcat KO Caco2 cells, taking advantage of the unique characteristics of each line.

### Junctional defects in αE-catenin-deleted cells

We next examined how cell junctions were affected by αE-catenin removal. As previously reported (Watabe-Uchida et al., 1998), DLD1 cells organized a typical AJC network, whereas αEcat KO DLD1 cells did not, showing dotted distribution of E-cadherin and tight junction (TJ) proteins at their cell-cell boundaries (Fig. S2A). These αEcat KO DLD1 cells tended to round up, preventing us from observing their peripheral structures closely. For detailed microscopic analysis, therefore, we mainly used Caco2 cells that show flatter morphology. In a migrating sheet of wild-type Caco2 cells, cells located at the front-most and sub-front regions, defined as marginal and sub-marginal cells, respectively, were flatter than more internal cells (Fig. 2A). In any region of the sheet, their AJs were characterized by linear distribution of E-cadherin and associated actin cables along the apical cell-cell boundaries. Ecadherin also distributed to lateral cell-cell contacts (LCs), which were generally slanted toward either side of the junction, exhibiting a strand-like or dotted pattern (Fig. 2B).

**Figure 2.**
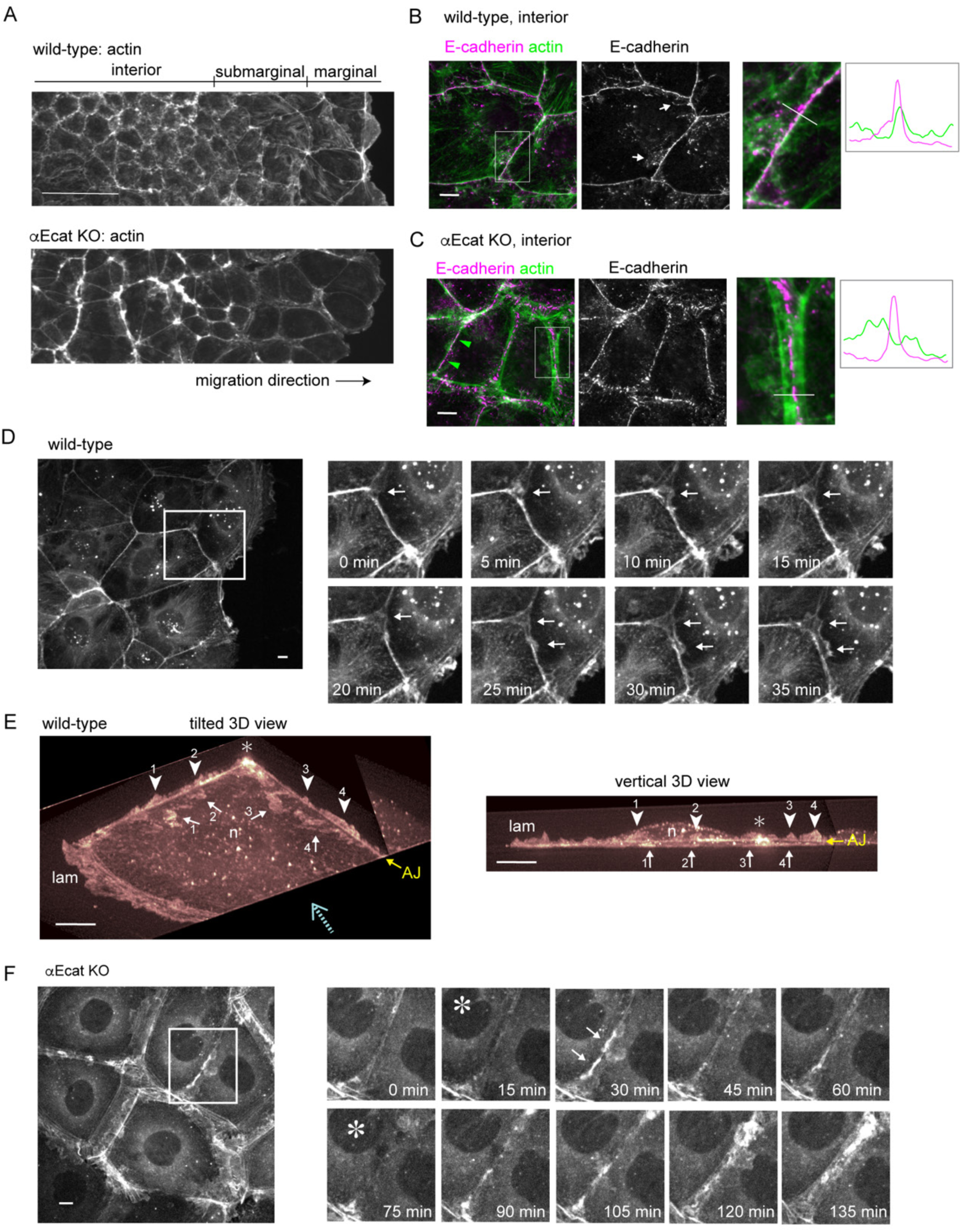
Actin assembly and protrusion formation at junctions. (**A**) Low-magnification view of a wild-type (A) or αEcat KO (B) Caco2 cell sheet, which is engaging in woundhealing movement. Stained for actin. (**B, C**) Co-staining for E-cadherin and actin in wild-type (B) and αEcat KO (C) Caco2 cells. Fluorescence signals in the boxed regions, which are enlarged at the right, were scanned along the white bar. White arrows indicate examples of Ecadherin-positive LCs. Green arrowheads point to a closed junction. (**D**) Montage of Video 3. Time-lapse images of the boxed region are shown. Arrows indicate emergence of protrusions. (**E**) LLSM images of a marginal wild-type Caco2 cell expressing Lifeact-RFP are displayed in 3D and viewed from two distinct angles. These were picked out from the images of Video 4, which had been edited to observe from desired angles. Arrows indicate protrusions that crawl under the cell shown here, and arrow heads point at another form of protrusions that grow upward. Numbers indicate identical structures at the left and right images. ‘AJ’ shows the position of this junction, which is 1.5 to 2.0 μm high above the bottom of the cell. Dotted arrow shows an approximate direction for the vertical observation at the right panel. lam, lamellipodium; n, the position of nucleus; asterisk, the position where two bicellular junctions merge. The images were cropped three-dimensionally for better visualization of vertical protrusions. (**F**) Montage of Video 9. Time-lapse images of the boxed region are shown. Arrows indicate representative protrusions whose appearance is oscillating. Asterisks, images in which protrusions are temporally silent. Scale bars, 100 μm for A; 10 μm for B-F.

αEcat KO Caco2 cells retained intercellular contacts but exhibited various defects in the junctions. Actin cables running along AJs were split into parallel lines at cell-cell contact sites, resulting in formation of a gap between them (Fig. 2C). E-cadherin was detected within this gap in a fragmentary pattern. The gaps contained an abundance of amorphous F-actin networks, but E-cadherin did not show any close co-localization with them (Fig. 2C), agreeing with the model that cadherins are normally linked to F-actin via αE-catenin (Takeichi, 2014). E-cadherin partners, β- and p120-catenin, showed distributions nearly identical to that of E-cadherin, as expected (Fig. S2B). Desmosomal proteins were also concentrated within these gaps (Fig. S2C). These observations suggest that, in the absence of αE-catenin, E-cadherin still helps bind the plasma membranes together, along with desmosomes, despite its failure to associate with F-actin. On the other hand, TJ proteins were localized along the split actin cables (Figs. S2D and S2E). These changes, induced by αEcatenin loss, were observed throughout a cell sheet (Fig. 2A). Importantly, the actin-cable splitting did not always occur in the αEcat KO Caco2 cell sheets, that is, a given single cell often had non-split junctions, too, at its borders with other neighbors (Fig. 2C). In these junctions, not only E-cadherin but also desmosomal and TJ proteins accumulated together (Figs. S2C and S2D). For convenience, we hereafter refer to the junctions where actin cables split and did not split as ‘open’ and ‘closed’ junctions, respectively.

### Cryptic lamellipodia form under the control of AJs

To investigate how the cadherin-mediated junctions or AJs control epithelial sheet migration, we observed actin dynamics using wild-type or αEcat KO Caco2 cells that were stably transfected with Lifeact-RFP, an F-actin binding peptide (Riedl et al., 2008), since actin plays a central role in cell migration. Live imaging of a wild-type cell sheet, which was undergoing wound healing, showed that all migrating cells were firmly connected together at the level of AJs, as assessed by the stable appearance of AJ-associated actin networks (Video 3). Closer observation, however, indicated that fan-shaped protrusions sporadically emerged from the AJ-associated actin cables, which likely correspond to c-lamellipodia (Farooqui and Fenteany, 2005). Such protrusions occurred at the junctions of marginal and sub-marginal cells, but less clearly occurred in interior cells that are taller and migrating more slowly than the front cells, as noted before (Farooqui and Fenteany, 2005). Frequently, protrusions initially arose from a multicellular junction, then extended toward the bicellular sites of the junction (Fig. 2D). For more detailed analysis of protrusion formation, we used lattice lightsheet microscopy (LLSM). Three-dimensional imaging by LLSM unexpectedly revealed that there are two forms of protrusion (Video 4 and Fig. 2E). One was a flat projection (with a thickness less than one μm) that invade underneath the adjoining cell, which structurally corresponds to c-lamellipodium. The other form of protrusion arose upward from the AJ zone, whose orientation was similar to that of ruffling membranes which occur at the front edges (or lamellipodia) of migrating cells. Whether these upward protrusions grew from both sides of the junction or only from a single side was not clear. These observations indicated that AJs function not only in tying cells together, but also as a site to generate c-lamellipodia and other dynamic protrusions.

We additionally observed how actin dynamics changes during AJ formation, and how the AJs enable cells to migrate. To this end, we took movies of singly isolated cells labeled with Lifeact-RFP, and their descendants. Isolated wild-type Caco2 cells tended to display a disc-like shape, generating lamellipodia all along the cell periphery (Video 5). When a cell divided into a pair, the descendants promptly organized AJ-associated actin cables between them. Such pairs of cells often showed rotating movement, but never displayed migration. After further divisions, they came to form a multicellular colony. A marginal cell in the colony, which was surrounded by two to three neighbors, sporadically began migration using its lamellipodia that formed at the free edges (Video 6). Thus, the AJ begins to control cell motility at the two-cell stage, but it requires more cells to organize a polarized cell sheet in order to conduct directed migration.

In the case of αEcat KO Caco2 sheets, moving cells exhibited lamellipodia-like protrusions everywhere at their periphery (Video 7). AJ-like actin cables were detectable, but not stable. The behavior of isolated αEcat KO Caco2 cells was indistinguishable from that of isolated wild-type cells. However, when they formed a pair, vigorous membrane ruffling continued even at the cell-cell contact sites (Video 8), and this feature persisted after further division of the cells (Video 9). Movies also revealed that, in colonies of αEcat KO Caco2, the open and closed junctions dynamically converted from one to the other, and the junctional closure resulted in a temporary suppression of membrane ruffling (Fig. 2F), which implies that the observed membrane dynamics can be controlled by simple mechanical cell-cell contacts in the absence of αE-catenin. Thus, in the absence of αE-catenin, it seems that cells are unable to control c-lamellipodia formation, but exhibiting an altered form of contact-dependent regulation of protrusion formation.

### WAVE complex is required for junctional membrane protrusion

We began to explore how AJs generate c-lamellipodia. Since the WAVE regulatory complex (WRC) is known as a major regulator of actin assembly in lamellipodia (Takenawa and Suetsugu, 2007), we examined their potential contribution to c-lamellipodia formation using Caco2 cells. To determine subcellular distribution of WRC components, we initially observed three of them, Abi1, WAVE2 and Nap1, finding that they were essentially identical in distribution (Figs. S3A and S3B). In the following experiments, we observed a representative one out of the three, unless otherwise noted. Consistent with previous reports (Han et al., 2014; Nishimura et al., 2016; Verma et al., 2012; Yamazaki et al., 2007), Abi1 was localized to AJs, overlapping with E-cadherin, in interior cells of wild-type cell sheets (Fig. 3A left). In marginal and sub-marginal cells whose AJs show protrusions, however, Abi1 was detected not only along the junctions, but also at the edges of the protrusions (Fig. 3A right). The frequency of the Abi1-positive protrusions considerably varied from junction to junction, as expected from their dynamic nature observed in movies. E-cadherin did not localize at the edges of such protrusions, although it was detected inside the larger protrusions in particular.

**Figure 3.**
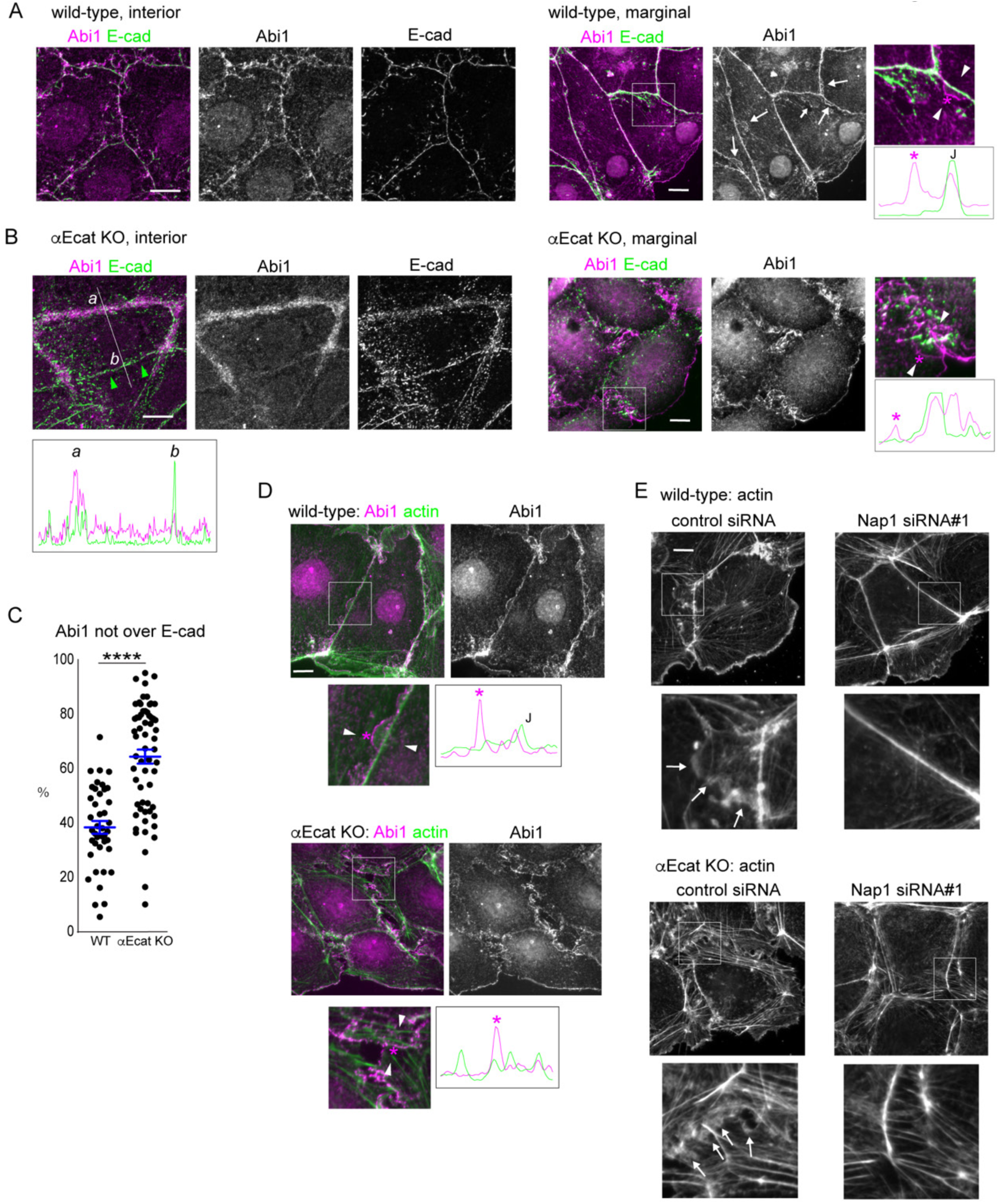
WAVE complex at junctions. (**A, B**) Co-immunostaining for Abi1 and E-cadherin in wild-type (A) and αEcat KO (B) Caco2 cells. Immunofluorescence signals in the boxed regions, which are enlarged at the right, were scanned along the line (not shown) drawn between the white arrowheads. Asterisk in the image indicates a position roughly corresponding to the peak labeled with the same symbol on the scan, throughout the figures. J, the position of cell junction. In B, immunofluorescence signals were also scanned along the line marked with *a* and *b*. Larger arrows point to Abi-positive protrusions; small arrow, a protrusion having E-cadherin. Green arrowheads point to a closed junction. (**C**) Immunofluorescence signals for Abi1 that overlap or do not overlap those for E-cadherin were measured using interior cells, and the ratio of the non-overlapping to the total signals was plotted. For measurement, a few points in each of the bi-cellular junctions were randomly selected, using four wild-type and three αE-cat KO cells. ****p<0.0001 (t-test). (**D**) Co-immunostaining for Abi1 and actin in wild-type or αEcat KO Caco2 cells. Immunofluorescence signals were scanned as explained in A. Marginal and sub-marginal cells are shown. J, the position of cell junction. (**E**) Effect of siRNA-mediated Nap1 depletion on actin assembly. The boxed areas are enlarged, being placed under each panel. Arrows indicate protrusions. Marginal and sub-marginal cells are shown. Scale bars, 10 μm.

In αEcat KO Caco2 cell sheets, the open junctions in interior cells were filled with Abi1-bearing amorphous structures (Fig. 3B left), and these were detected as fan-shaped membranes in marginal and submarginal cells (Fig. 3B right). Quantitative measurement indicated that Abi1-positive membranes, which do not overlap with E-cadherin, were increased at those junctions (Fig. 3C). In the closed junctions, on the other hand, the relative level of Abi1 was reduced (Fig. 3B left scan), according with the observation that membrane ruffling was suppressed at these sites. DLD1 cells also showed a similar overlapping of Abi1 with E-cadherin at wild-type junctions, and a separation of Abi1-postive membranes from Ecadherin in the absence of αE-catenin (Fig. S3C). To summarize, Abi1 localizes at Ecadherin-positive AJs, but it becomes distributed also to protrusions when they form, and the latter fraction of Abi is increased at the open junction of αEcat KO cells.

Double-staining for Abi1 and actin confirmed that they overlap with one another at both protrusions and junctions in wild-type cells (Fig 3D upper). Live imaging of Nap1-GFP introduced into Lifeact-RFP transfectants showed that Nap1/actin double-positive membranes dynamically protruded from AJ regions (Video 10), suggesting that WRC redistributes from AJ to the protrusion during its formation. In αEcat KO Caco2, Abi1-positive membranes lost any defined relation with particular actin cables (Fig. 3D lower), preventing us from identifying the origin of their formation.

To confirm the role of WRC in c-lamellipodia formation, we depleted Nap1 in Caco2 cells using siRNAs (Fig. S3D). Nap1 depletion resulted in disappearance of Abi1 from junctions (Fig. S3B), suggesting that WRC was disorganized there, and this treatment eliminated actin-positive protrusions that are associated with AJs (Fig. 3E top). The open junctions in αEcat KO Caco2 cells also lost fan-shaped protrusions, becoming filled with only with fibrous actins (Fig. 3E bottom). Thus, WRC is important for c-lamellipodium formation.

### Arp2/3 complex cooperates with WRC for protrusion formation

A major function of WRC is to activate the Arp2/3 complex (Arp2/3) which mediates actin nucleation (Krause and Gautreau, 2014; Rotty et al., 2013), prompting us to test whether Arp2/3 is also involved in c-lamellipodia formation. Immunostaining for p34/ARPC2, a subunit of this complex, showed that it localized along AJs (Fig. 4A), consistent with previous observations (Kovacs et al., 2002; Verma et al., 2012; Verma et al., 2004). This p34/ARPC2 overlapped with Abi1, E-cadherin and associated actin cables, although it did not distribute to LCs, unlike WRC. p34/ARPC2 was also detectable in actin-positive protrusions, distributing more diffusely than Abi1. In αEcat KO Caco2 cells, similar overlapping of p34/ARPC2 and Abi1 was seen along protrusions in the open junctions (Fig. 4B). At basal regions of these protrusions, p34/ARPC2 and Abi1 irregularly aggregated together with actin clusters, instead of localizing at AJs. Notably, E-cadherin lost its overlapping with p34/ARPC2 in these cells (Fig. 4B bottom). Thus, the co-distribution of these molecules at AJs was disorganized in the absence of αE-catenin. The expression levels of WRC components and p34/ARPC2 were not affected by αE-catenin loss (Fig. S4A).

**Figure 4.**
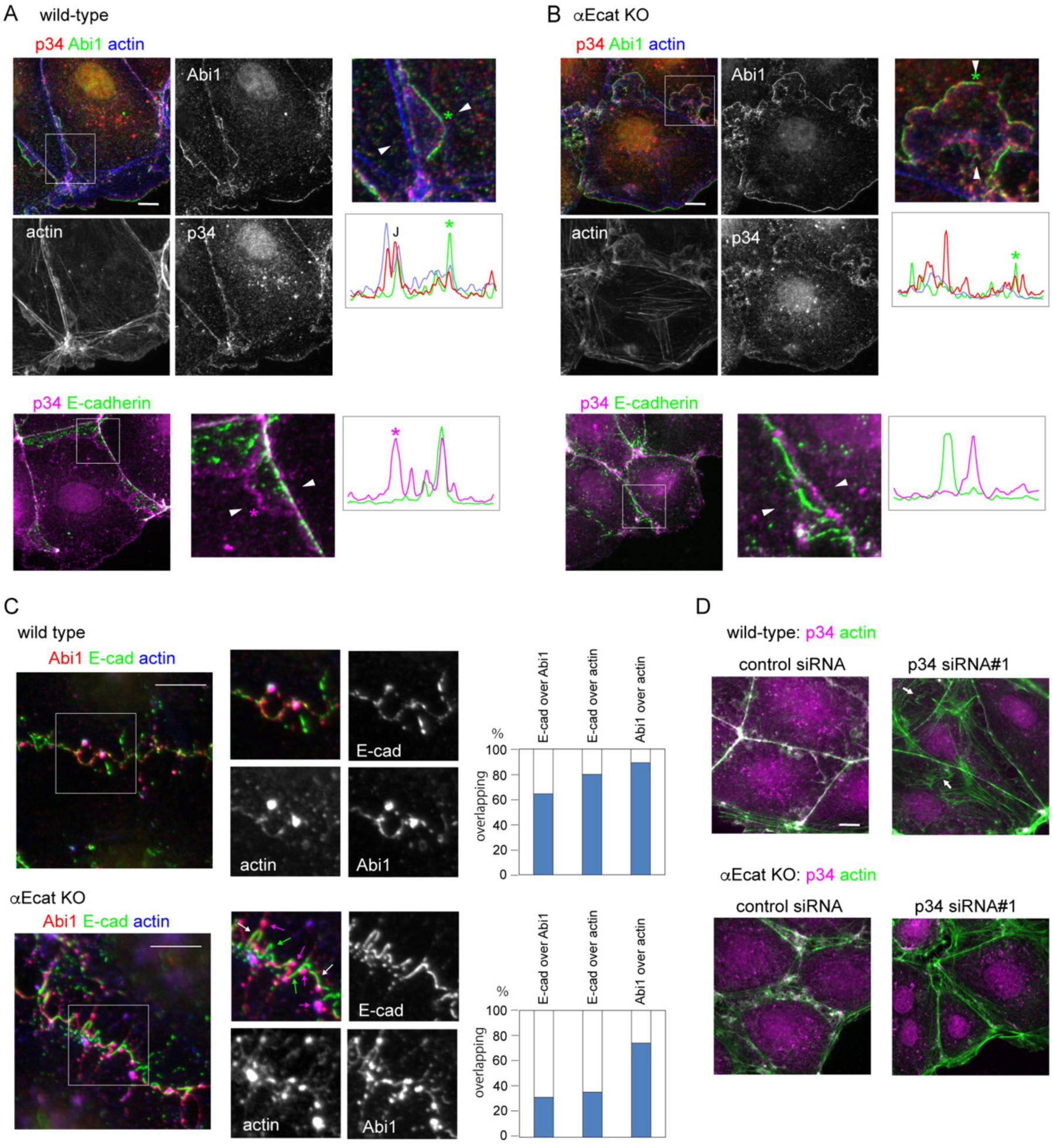
Arp2/3 complex at junctions and its interaction with WRC and F-actin. (**A, B**) Co-immunostaining for p34/ARPC2 (p34), Abi1 and actin (top); and for p34/ARPC2 and Ecadherin (bottom) in wild-type (A) or αEcat KO (B) Caco2 cells. J, the position of junction. Marginal or marginal plus sub-marginal cells are shown throughout this figure. (**C**) Coimmunostaining for Abi1, E-cadherin and actin in wild-type or αE-cat KO Caco2 cells, which were incubated with 10 μM latrunculin A for 60 min. The boxed regions are enlarged. The overlapping ratios of these molecules were quantified. Magenta arrows, Abi1–actin coclusters; white arrows, Abi1–E-cadherin–actin co-clusters; green arrows, E-cadherin fragments that do not overlap with other molecules. (**D**) Effect of p34/ARPC2 siRNA treatment on actin assembly. Arrows point to actin-positive structures that are not closely associated with the junctions. Scale bars, 10 μm.

Previous studies suggested that WRC, Arp2/3 and E-cadherin physically interact with each other via cortactin (Han et al., 2014; Kovacs et al., 2002). To gain further insights into how these molecules associate with AJs, we examined the effect of actin disturbance on their distribution, because all of them directly or indirectly interact with F-actin. Treatment of wild-type Caco2 cells with an actin polymerization inhibitor, latrunculin A, resulted in clustering of actin, leaving strand-like actin linkages between the clusters (Figs. 4C). Abiwell co-clustered with these reorganized actin molecules, and E-cadherin also showed some level of overlapping with actin or Abi1, even alter the sever disturbance of actin assembly. In αEcat KO cells, however, although Abi1 still overlapped with actin clusters, E-cadherin became less co-aggregative with Abi1 or actin. p34/ARPC2 behaved in a way similar to Abi1 in these latrunculin A-treated cells. These observations suggest that, in normal AJs, actin filaments serve as a core for E-cadherin, WRC and Arp2/3 to assemble together, and in the absence of αE-catenin, E-cadherin leaves from this assemblage, due to the loss of its actin-binding partner.

To confirm the role of Arp2/3 in c-lamellipodia formation, we treated cells with siRNA for p34/ARPC2 (Fig. S4B). This treatment caused regression of actin-positive protrusions in both wild-type and αEcat KO cells, although some irregular actin-positive structures, which are separated from AJs, remained in wild-type cells (Fig. 4D). We also examined the effect of an Arp2/3 inhibitor, CK666, using Lifeact-RFP–expressing Caco2 cells. Movies showed that this inhibitor suppressed dynamic protrusion of fan-shaped membranes, although it permitted extension of some flat structures (Video 11, right). Since the WRC-Arp2/3 signaling is regulated by Rac1 GTPase (Chen et al., 2017), we also examined the effect of an Rac inhibitor, EHT 1864 (Shutes et al., 2007). In the treated wild-type or αEcat KO cells, Abi1-positive protrusions disappeared, and both Abi1 and p34/ARPC2 came to accumulate along AJ-associated actin filaments in wild-type cells, or irregular actin aggregates in αEcat KO cells (Fig. S4C). These observations confirmed that the Rac1-dependent WRC-Arp2/3 system is important for c-lamellipodia formation, and also suggested that AJs can hold non-activated WRC and Arp2/3 components.

### WRC-Arp2/3 system is required for collective migration of epithelial cells

We then tested whether c-lamellipodia formation is really required for the collective migration of Caco2 cells. Prior to this test, we collected data regarding whether Nap1 or p34/ARPC2 depletion affected cell junctions. Depletion of each molecule did not affect the expression of the other molecule, nor E-cadherin, except that Nap1 depletion also removed its partner, WAVE2 (Fig. S5A), as was found for Abi1. Immunocytological analysis, however, showed that depletion of Nap1 or p34/ARPC2 caused reduction of the other at the junctions, suggesting that their junctional recruitment is inter-dependent (Figs. 5A and 5B). Notably, the area of E-cadherin-positive LCs dramatically increased in these cells (Figs. 5A and 5C), which suggests that AJ-derived protrusions are converted to simple adhesive structures when these actin regulators were silenced.

Next, we examined the migration of Caco2 cells in which Nap1 or p34/ARPC2 was depleted, by wound healing methods, and found that their migration was delayed (Fig. 5D). In this experiment, however, depletion of the actin regulators must have also affected the functions of genuine lamellipodia formed by the leader cells. To examine specifically the role of Napor p34/ARPC2 in the migration of follower cells, we prepared Lifeact-RFP–transfected Caco2 cells in which these molecules were depleted, and mixed them with non-labeled wildtype cells in a 1-to-1 ratio, subjecting the cell mixture to wound healing assays. The results showed that Nap1 or p34/ARPC2-depleted cells were left behind during the movement of cell sheets (Figs. 5E and 5F), confirming that the WRC-Arp2/3 system is required for the migration of follower cells. In these experiments, cell proliferation rate did not differ between control, Nap1, and p34/ARPC2 siRNA-treated cells, in which the percentage of mitotic cells in culture were 7.69±0.70 (n=6), 8.54±0.25 (n=6, p=0.3 vs control), and 7.07±0.36 (n=6, p=0.6 vs control), respectively.

**Figure 5.**
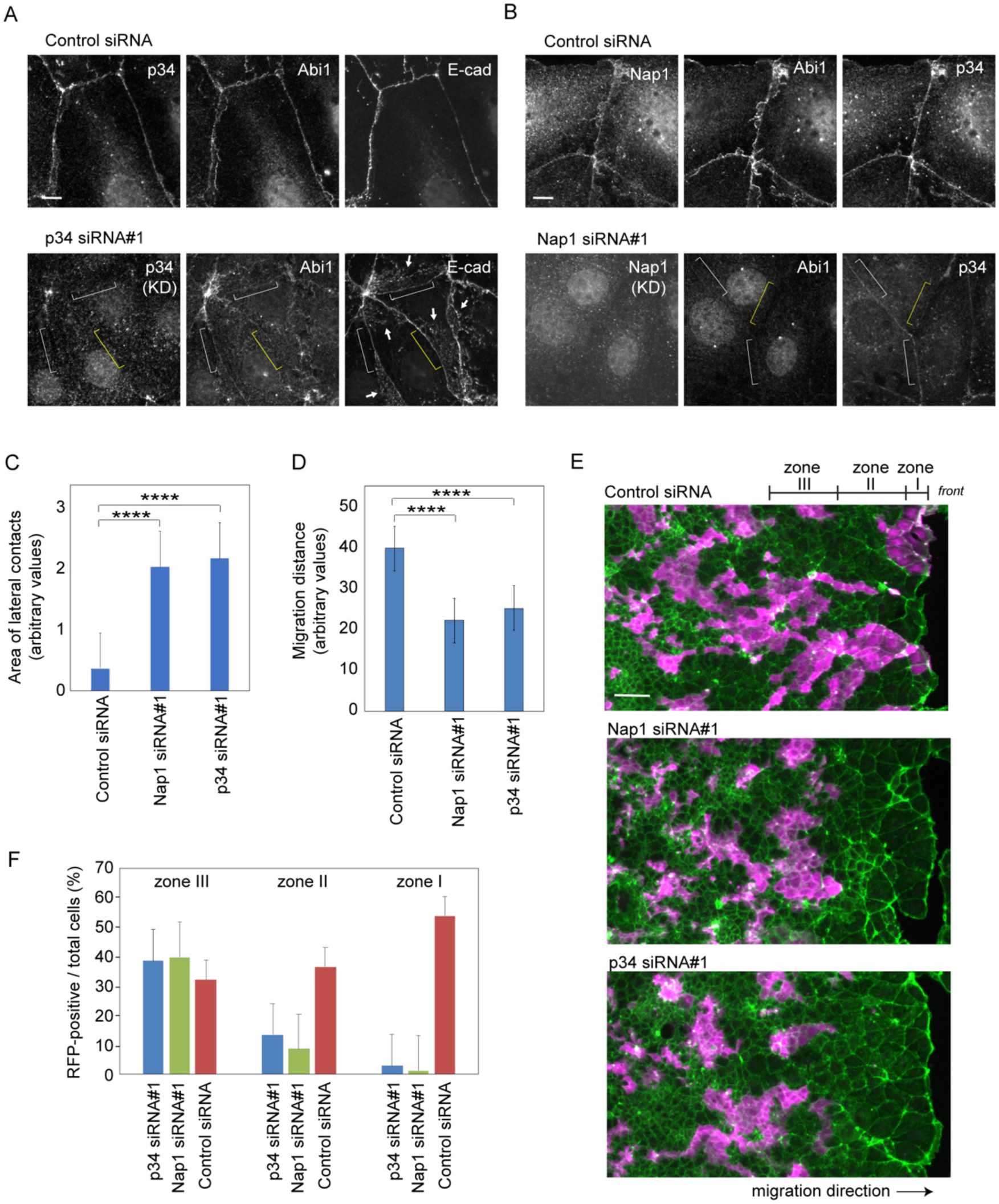
Effects of p34/ARPC2 or Nap1 depletion on collective cell migration. (**A, B**) p34/ARPC2 (A) or Nap1(B) was depleted using siRNAs, followed by co-immunostaining for the indicated molecules. Yellow brackets indicate a region of cell junction where the siRNAtargeted molecule is undetectable in immunostaining. White brackets indicate a region where p34/ARPC2 or the Nap1 partner Abi is still faintly visible. Arrows point to LCs that are identified by E-cadherin distribution. Marginal and sub-marginal cells are shown. (**C**) Areas of E-cadherin-positive LCs in marginal and sub-marginal cells. The entire area of LC expanding below from each bilateral AJ was measured and compared. 31 junctions in control cells, and 23 junctions in Nap1 or p34/ARPC2 siRNA-treated cells were analyzed. ****p<0.0001 (t-test). (**D**) Migration distance of the marginal cells during 24 hr in wound healing cultures. Values were obtained using four and six independent areas for control and Nap1 or p34/ARPC2 siRNA-treated cells, respectively. ****p<0.0001 (t-test). (**E, F**) Sheets of wild-type Caco2 cells that are mixed in a 1-to-1 ratio with those expressing Lifeact-RFP in which Nap1 or p34 p34/ARPC2 has been depleted with siRNAs. Magenta, Lifeact-RFP; green, actin. Photographed at 48 hr after wounding. In F, the ratio of siRNA-treated (Lifeact-RFP-labeled) cells to the total cells in the three tandem zones of a culture was measured. Zones I, II and III roughly correspond to a row of marginal cells, rows of sub-marginal cells, and an anterior group of interior cells, which spans 58, 188 and 188 μm in width, respectively. We analyzed 11, 13 and 5 images for control, Nap1 and p34 p34/ARPCsiRNA-treated cultures, respectively. Scale bars, 10 μm A and B; 100 μm in E.

### AJ disruption induces myosin II activation

Next, we investigated whether there is any mechanism to ‘control’ c-lamellipodia formation, as it occurred in a non-persistent way in wild-type cells, compared with the constant production of them in αEcat KO cells. As mentioned already, a hot spot to generate membrane protrusion was multicellular junctions where multiple bicellular junctions meet (Fig. 2D), and the multicellular junctions are thought to be prone to disruption, as they receive tensile force from the bicellular junctions (Higashi and Miller, 2017; Stephenson et al., 2019). Such force is generated by contraction of junction-associated actomyosin, and therefore we examined whether there is any relation between myosin II activation and clamellipodia formation. Myosin II is activated by Rho kinase/ROCK via phosphorylation of myosin regulatory light chain 2 (MLC2) (Vicente-Manzanares et al., 2009). Immunostaining for Thr18/Ser19-phosphorylated MLC2 (ppMLC2) showed that, in wild-type Caco2 cells, ppMLC2 was detected along some junctions of marginal and sub-marginal cells, but not in those of interior cells (Fig. 6A, leftmost), suggesting that cells in the marginal and submarginal zones receive higher tension than interior cells, as myosin II activation is known to be a tension-sensitive process (Fernandez-Gonzalez et al., 2009). However, ppMLC2 localization did not perfectly correlate with the abundance of Abi1-posive membranes at AJs.

**Figure 6.**
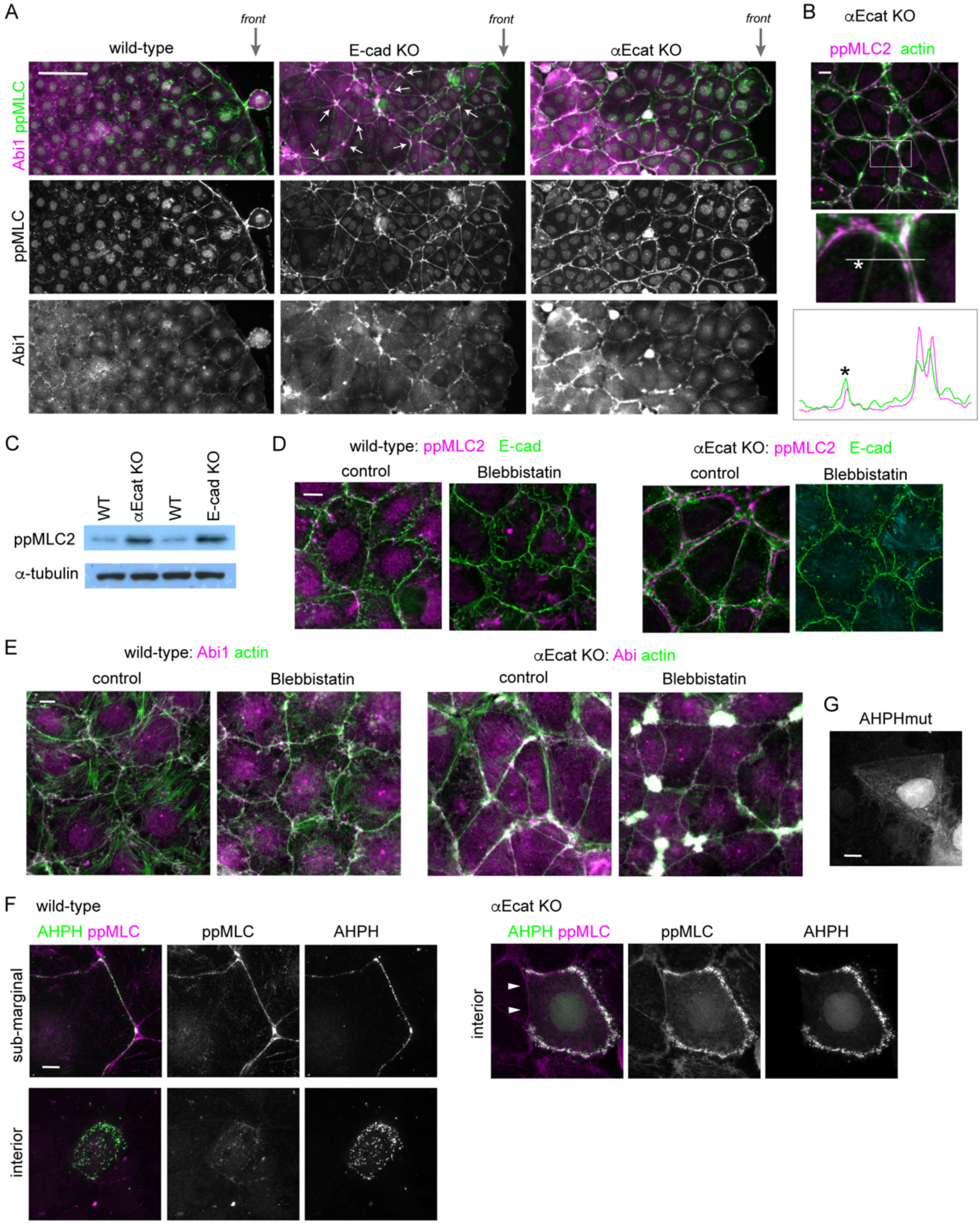
Junction disruption induces myosin II activation. (**A**) Low-magnification view of the indicated Caco2 cell sheets, which are engaging in wound healing, coimmunostained for ppMLC2 and Abi1. ppMLC2 and Abi are co-condensed in approximately 40% of multicellular junctions in E-cad KO cells, examples of which are indicated by arrows, whereas the junctions in wild-type cells do not show such features. *Front* indicates the front regions of the migrating cell sheet. (**B**) Co-immunostaining for ppMLC2 and actin in αE-cat KO Caco2 cells. The boxed area was enlarged, and ppMLC2 and actin were scanned along the line indicated. (**C**) Western blots for ppMLC2 in the indicated Caco2 lines. (**D, E**) Wildtype or αEcat KO Caco2 cells were incubated with 10 μM blebbistatin for 6 hr, and double-immunostained for ppMLC2 and E-cadherin (D), and Abi1 and actin (E). Cells located around the inner to sub-marginal zones were photographed. (**F**) Distribution of the Rho-GTP localization sensor GFP-AHPH. Wild-type or αEcat KO Caco2 cells were transiently transfected with GFP-AHPH, and then double-immunostained for GFP and ppMLC2. Arrowheads indicate a closed junction in an αE-cat KO Caco2 cell. (**G**) A wild-type cell transiently transfected with a function-deficient mutant of AHPH (AHPHmut). Scale bars, 100 μm for A; 10 μm for B, D to G.

To further test the potential involvement of myosin II in c-lamellipodia regulation, we introduced *CDH1*-deleted Caco2 cells (E-cad KO Caco2 cells) to the experiments (Fig. S1B). These cells still maintain epithelial sheets, in which p120-catenin, a cadherin partner, normally localized along cell junctions, although its level was greatly reduced (Fig. S5B), suggesting that they sustain AJs, likely due to the presence of some classical cadherins other than E-cadherin. Consistently, TJ network and p34/ARPC2 distribution also looked normal (Fig. S5C). We used these cells as a model that has normal-looking junctions but reduced adhesive molecules. Western blots showed that ppMLC2 level was increased in E-cad KO cells (Fig. 6C), and ppMLC2-positive cells extended to deeper regions of their sheets (Fig. 6A, middle), indicating that the junctions with less cadherins are more susceptible to myosin II activation. Then, we examined the distribution of Abi1 in these cells, and found that it was condensed at multicellular junctions, along with condensed ppMLC2, whereas Abi level at the bilateral junctions was comparable to that in wild-type cells. These findings support the idea that WRC-dependent protrusion occurs most easily at mechanically weak points of the junction. Furthermore, in αEcat KO Caco2 cells, ppMLC2 was detected throughout the cell layer along the split actin cables in the open junctions (Figs. 6A, rightmost), accompanied by an increase of the total ppMLC2 level (Fig. 6C). The closed junctions, however, exhibited lower levels of ppMLC2 (Fig. 6B). Thus, myosin II activation occurred in correlation with reduction of cadherins or junction disruption.

To confirm the impact of myosin II activation on junction or protrusion formation, we treated cells with blebbistatin, an inhibitor of myosin II (Straight et al., 2003). Although this treatment did not much affect wild-type junctions in terms of their integrity and Abi1 distribution, it induced closure of junctions in αEcat KO Caco2 cells, except at multicellular junctions where amorphous actin and Abi remained to cluster (Figs. 6D and 6E). At the closed bilateral junctions, the extent of Abi1-positive protrusions became comparable to those in wild-type junctions or even reduced (Fig. 6E). These results suggest that AJ disruption enhanced myosin II activation and the resultant contraction of actomyosin cables induces cell separation, leading to uncontrolled production of c-lamellipodia. However, it remains unclear whether myosin II activity controls c-lamellipodia formation in the normal situations.

Since myosin II is activated by RhoA and its effectors, we observed active RhoA localization. We transiently transfected Caco2 cells with a biosensor for active RhoA containing the RhoA-binding domain of anillin, GFP-AHPH (Tse et al., 2012), and found that, in wild-type cells, this probe was detected along some junctions of marginal or submarginal cells, but not of interior cells (Fig. 6F left). In the case of αEcat KO Caco2 cells, GFP-AHPH was detected in the open junctions even at interior portions of a cell sheet, whereas it never came to the closed junctions present in the same cell (Fig. 6F right). Importantly, these distributions of GFP-AHPH correlated with the ppMLC2 level at the junctions. The specificity of the GFP-AHPH probe employed in these experiments was confirmed using its mutated, non-functional version, which mostly diffuses in the cytoplasm (Fig. 6G). These results suggest that myosin II activation induced by junction disruption was probably mediated by RhoA activation at cell-cell contact areas.

### AJ disruption interferes with epithelial cell migration through myosin IIA activation

We finally tested whether myosin II-dependent AJ disruption was involved in the impaired migration of epithelial cells, using the DLD1 line, as migration of this line is more sensitive to AJ loss than that of Caco2 (Fig. 1B). Initially we checked whether myosin II activation is responsible for AJ disruption also in this cell line. We prepared GFP-tagged MLC2 mutants in which Thr18 and Ser19 were replaced with Ala (AA-MLC2) and Asp (DD-MLC2) to generate a constitutively inactive and active construct, respectively. When AA-MLC2 was introduced into αEcat KO DLD1 cells, it restored a normal-looking AJC, whereas DD-MLC2 expression had no such effects (Fig. 7A). We next explored which of myosin IIA and IIB heavy chains was important as the partner for ppMLC2 in the regulation of AJs. When *MHY9* (*NMMHC-IIA*) was deleted in αEcat KO DLD1 cells, AJC-like reorganization was partly recovered, whereas removal of *MHY10 (NMMHC-IIB)* had no clear effects (Fig. 7B), indicating that myosin IIA-based actomyosin contraction is more important in inducing junction disruption. Then, we tested whether these changes in myosin II affected cell migration using wound-healing assays, and found that AA-MLC2 expression in αEcat KO DLD1 cells promoted their wound healing more vigorously than DD-MLC2 expression (Fig. 7A). Likewise, deletion of myosin IIA, but not myosin IIB, promoted migration of αEcat KO DLD1 cells (Fig. 7B). These findings suggest that the molecular events that interferes with the collective migration of epithelial cells whose AJs have been disrupted include myosin IIA activation.

**Figure 7.**
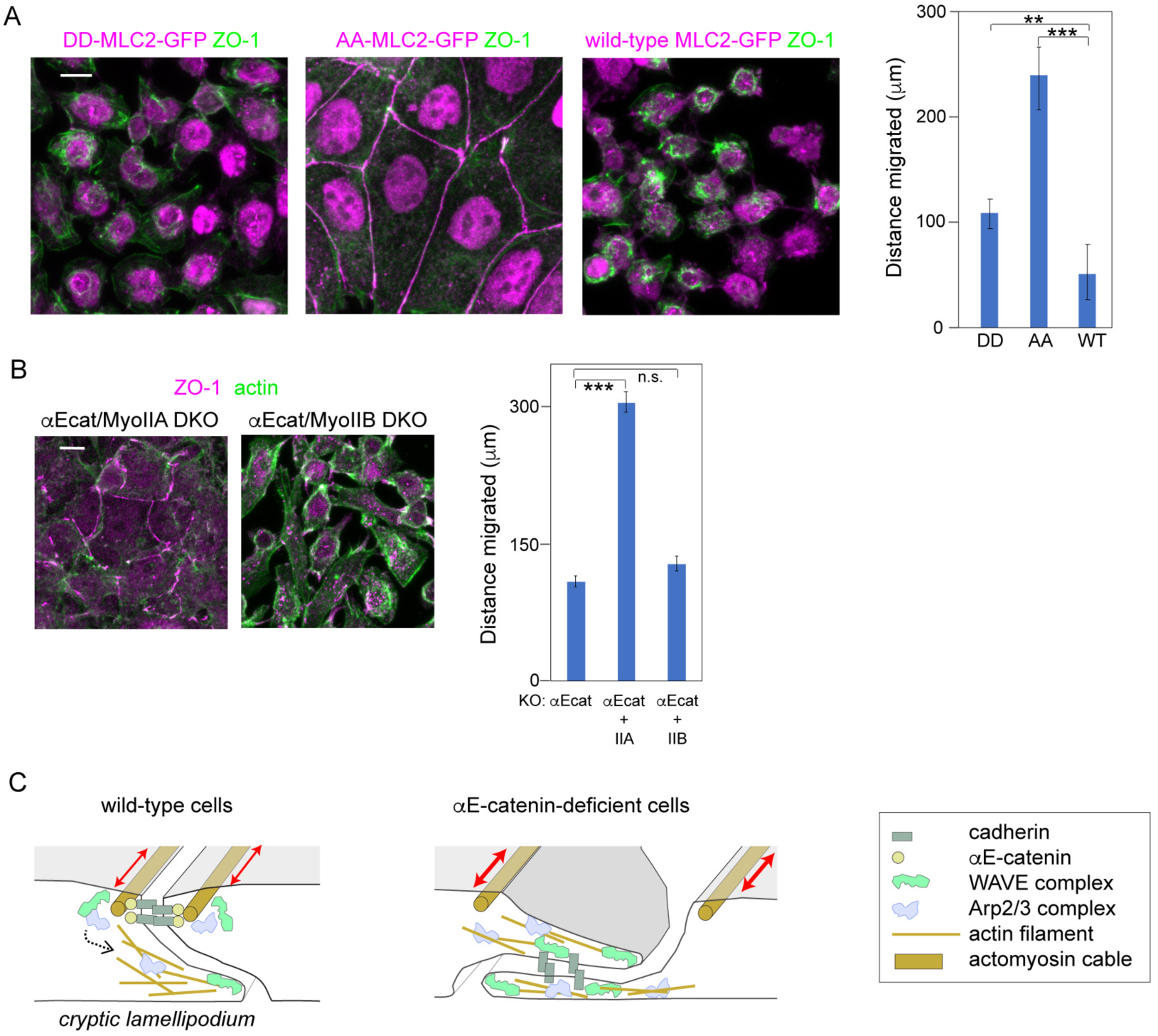
Tests for the role of myosin II activation in migration of αEcat KO DLDcells, and summary of the results. (**A**) Stable lines of αEcat KO DLD1 cells transfected with the indicated MLC2 mutants were immunostained for GFP and ZO-1. (**B**) Myosin IIA or IIB was deleted in αE-cat KO DLD1 cells, as shown in Fig. S1C. Cells were immunostained for ZO-1 and actin. Graphs, migration distance of cells at the wound edges 24 hrs after scratching of the culture. Data were analyzed as explained in the legend of Fig. 1B. (**C**) Illustrated summary of the results. In wild-type cells, WAVE and Arp2/3 components are anchored to AJ and are involved in c-lamellipodia formation when activated by undefined mechanisms (dotted arrow). In the open junction of αEcat KO cells, WAVE and Arp2/components appear to be constitutively active, without having any specific anchoring structures. Contraction of AJ-associated actomyosin cables (two-way arrows) causes AJ disruption in αEcat KO cells. Scale bars, 10 μm.

## Discussion

Mechanisms of cell migration have extensively been studied using various cell types (De Pascalis and Etienne-Manneville, 2017). The behavior of ‘isolated’ epithelial cells, however, has not so closely been observed except for restricted cell types. Our results showed that, when singly isolated, adenocarcinoma-derived epithelial cells as well as MDCK cells were unable to migrate by themselves at least in the studied conditions, as reported before (Desai et al., 2009). In these isolated cells and also in AJ-disrupted cells, lamellipodia (or clamellipodia) randomly emerged from multiple sites of the cell periphery, explaining why they are unable to move toward a particular direction. By contrast, in wild-type cell sheets, the leader cells were polarized to protrude lamellipodia only at the free edges, and, within the sheets, AJs restrain individual cells from free movement, resulting in well-ordered migration of the cells.

For the entire epithelial sheet to move, however, not only the leader cells but also follower cells must actively move. Actually, in a moving cell sheet, individual cells are known to form protrusions at their basal sides (Barlan et al., 2017; Farooqui and Fenteany, 2005; Krndija et al., 2019; Squarr et al., 2016). Our present study has now provided evidence for the importance of such protrusions or c-lamellipodia for adenocarcinoma cells to move as a collective. c-lamellipodia were generated as dynamic membrane protrusions from AJs, and this process depended on WRC and Arp2/3, whose components were localized along AJs. Given the well-known function of these molecules at lamellipodia, we can infer that junctional WRC–Arp2/3 serves for actin nucleation so as to induce c-lamellipodia formation (Fig. 7C left). c-Lamellipodia were morphologically similar to slanted LCs, as larger clamellipodia actually contained E-cadherin. Of note, silencing of WRC or Arp2/3 resulted in an increase of E-cadherin-positive LCs, along with suppression of dynamic protrusions. This suggests that the WRC–Arp2/3 system normally functions to sustain c-lamellipodia formation, but, when this system is inactivated, the protrusions are converted to stable LCs. It remains to be determined whether the AJ-dependent mechanism uncovered here using ‘thin’ epithelial cells also works for protrusion formation in cuboidal or columnar epithelial cells, where AJs are located quite distant from the basal sides. LLSM analysis revealed that protrusions also grow upward at AJs, as seen during the retraction of ruffling membranes at the leading edge of migrating cells. This suggests that similar molecular events occur at both AJs and free edges. The biological function of upward protrusions, however, remains to be investigated in the future. Previous studies indicated that WRC and Arp2/3 are necessary for the integrity of zonula adherens (Verma et al., 2012), suggesting the possibility that this actin-regulating system might differently take part in junction dynamics in moving and stationary cells.

Since c-lamellipodia formation is an intermittent process, there could be a mechanism to control WRC–Arp2/3 activity during cell migration. In the case of αEcat KO cells, clamellipodia freely grew at open junctions, whereas they disappeared at closed junctions. In the latter junctions, Abi1 level was reduced, likely because AJs as a center for WRC recruitment were absent, explaining why protrusion growth halted here. In normal cells, however, WRC was always detectable along AJs, nevertheless, extension and retraction of protrusions were repeated. This suggests that the activity of WRC and Arp2/3 held by AJs is controlled by some sporadic signals. We suspect that mechanical signals might play a role in this hypothetical process, as our results indicated that mechanical weakening or disruption of AJs enhances protrusion formation, that is, in E-cadherin–deleted cells and even in wild-type cells, Abi-positive membranes tended to grow from multicellular junctions, where cell-cell contacts are thought to be prone to disruption (Higashi and Miller, 2017). Our results also suggested that WRC and Arp2/3 bound to AJs are not necessarily active, as they stayed in AJs even when their upstream regulator Rac1 was inhibited. Based on these observations, we propose that mechanical changes in AJs may trigger activation of the WRC–Arp2/3 system. Such changes could be brought about by tensile force that is exerted to AJs. Actually, marginal and submarginal cells appeared to receive such force, as myosin II activation, which is known to be a tension-sensitive process (Fernandez-Gonzalez et al., 2009), was observed in their junctions. In the open junctions of αEcat KO cells, on the other hand, WRC and Arp2/3 were apparently not anchored to any special structures except for their target, F-actin, and this might cause a constitutive activation of these actin regulators and in turn uncontrolled protrusion growth (Fig. 7C right), as seen in the lamellipodia of free cell edges.

We demonstrated that myosin II activation played a crucial role in disrupting AJs under the αE-catenin-free condition. Inhibition of myosin II with inhibitors or MLC2 mutations led to restoration of cell-cell contacts in αEcat KO cells. Furthermore, active RhoA was detected only along the disrupted junctions, where myosin II was also activated. These observations suggest the presence of a signaling pathway that αE-catenin loss triggers activation of RhoA, and this in turn leads to MLC2 activation. The resultant contraction of AJ-associated actomyosin cables was, thus, apparently the primary cause of AJ disruption. How RhoA– MLC2 signals are activated by AJ loss remains to be elucidated. On the other hand, in the normal cadherin-bearing junctions, it appears that myosin II activation *per se* is not directly involved in WRC-Arp2/3 regulation, as the localization of ppMLC2 did not correlate with Abi1 level in wild-type cells. It is of note that junctional myosin II activation, which depends on cadherin reduction, was also observed *in vivo* (Hashimoto and Munro, 2019), suggesting that such mechanism might be used as a physiological tool for tissue reorganization.

Previous studies on the contact-inhibition of cell locomotion suggested that the motility of non-epithelial cells, such as neural crest and glioma cells, is inhibited through cadherin-mediated cell-cell adhesion (Hayashi et al., 2014; Theveneau et al., 2010). However, the contact-dependent control of cell motility appears to be more complicated in epithelial cells, as they keep producing c-lamellipodia at the cell-cell contact sites. Our results suggest that the epithelial AJ has a dual role: it works for stable linking of cells on the one hand, but on the other hand they work as a site to generate motile apparatus. Importantly, while WRC and Arp2/3 components accumulate at AJs in epithelial cells, these actin regulators are excluded from the cell-cell contact sites of other cells such as glioma, unless special mechanisms operate (Hayashi et al., 2014). This explains at least in part why different cell types differently respond to cell-cell contacts in motility regulation.

Cell migration and dissemination play a crucial role in cancer invasion and metastasis. Mechanisms for how tumor cells relocate themselves is a topic of controversy. E-cadherin or associated proteins have been found to be mutated in various types of carcinomas (Fanjul-Fernandez et al., 2013; Morrogh et al., 2012; Wang et al., 2014), and its loss promotes metastasis (Cai et al., 2014a). In other tumor cells, cadherin function is impaired even when it is expressed (Aono et al., 1999; Ito et al., 2017). Dysfunction of E-cadherin indeed causes tumor dissemination in certain cancers (Derksen et al., 2006; Nanki et al., 2018). On the other hand, cancer invasion often proceeds in the way of collective cell migration rather than free movement of dispersed cells (Friedl et al., 2012; Pandya et al., 2017), implying that invading tumor cells do not always lose cell-cell adhesion molecules. Recent studies showed that ductal carcinoma metastasis is driven by E-cadherin-positive cells but not by negative cells in a mouse model (Padmanaban et al., 2019). Our *in vitro* observations reported here would provide a clue to our deeper understanding of how E-cadherin or AJs are really involved in the invasive behavior of carcinoma cells.

## Materials and Methods

### Cell cultures and wound healing assay

DLD1 (a gift from Shintaro Suzuki, Kwansei Gakuin University), Caco2 (ATCC), MKN(JCRB Cell Bank), HT1080 (a gift from Kiyotoshi Sekiguchi, Osaka University), MDCK (a gift from Yasushi Daikuhara, Kagoshima University) and A549 (a gift from Varisa Pongrakhananon, Chulalongkorn University) were cultured in DME-M/Ham’s F12 medium (Wako) supplemented with 10% fetal calf serum (CCB Cell Culture Bioscience Cat#171012, lot 10L015) at 37°C, 5% CO_2_. For wound healing assays, cells were plated in a 6-well culture plate (IWAKI collagen type-I microplate 6well Cat#4810-010) and cultured for 24 h. A plastic pipette tip was used to draw a wound area across the center of the plate. Culture medium was then replaced with fresh medium. At 0, 12, 24, and 48 h in culture, cells were photographed. The distances of cell migration from original wound edge to the leading edge of the migrating cells were measured. For wound healing assays using siRNA-treated cells, cells were treated with siRNA for 6 hr, followed with replacement of the medium with a fresh one. After 48 hr, cells were dispersed by trypsinization, plated, and cultured overnight, before using for wound healing assays. When a mixed culture of siRNA-treated and non-treated cells was used, they were mixed in a 1-to-1 ratio before plating.

To detect mitotic cells in culture, cells were incubated for 24 h and then their monolayers were wounded. After another 24 hr, cells were fixed and immunostained for phosphorylated Histone H3 and DAPI (4′,6-Diamidino-2-phenylindole dihydrochloride). For detecting mitotic cells in siRNA-treated cultures, cells were pre-cultured overnight, then they were treated with siRNAs for 6 hr. After another 42-hr incubation with a fresh medium, cells were trypsinized, replated, and cultured overnight before measurement of mitotic cells.

### CRISPR/Cas9 plasmids

For CRISPR/Cas9-mediated knockout of genes, we used the pCGsapI vector developed by Takayuki Sakurai (Shinshu University) (Ozawa, 2018). The vector contains the *hCas9* gene under the control of the CAG promoter and a unique cloning site, *Sap*I site, for insertion of the guide RNA under the control of the U6 promoter. All synthetic oligonucleotides corresponding to the guide RNA and complementary chain, therefore, contain the adaptor sequence for *Sap*I. The following oligonucleotides were used to construct guide RNAs, in which the lowercase letters represent the adaptor sequences: For αE-catenin, accgGAAATGACTGCTGTCCATGCg and aaacGCATGGACAGCAGTCATTTCc; accgTCTGGCAGTTGAGAGACTGTg and aaacACAGTCTCTCAACTGCCAGAc; accgGAAGCGAGGCAACATGGTTCg and aaacGAACCATGTTGCCTCGCTTCc; accgGTCAGCCAAAATCAGCAACCg and aaacGGTTGCTGATTTTGGCTGACc. For E-cadherin, accgCCCTTGGAGCCGCAGCCTCTg and aaacAGAGGCTGCGGCTCCAAGGGc; accgGAGCCGGAGCCCTGCCACCCg and aaacGGGTGGCAGGGCTCCGGCTCc. For Myosin IIA, accgCCCGCCCAAGTTCTCCAAGGg and aaacCCTTGGAGAACTTGGGAGGGc. For Myosin IIB, accgCCCACCTAAGTTTTCCAAGGg and aaacCCTTGGAAAACTTAGGTGGGc. The vectors were introduced into cells together with the pCAG/bsr-7 vector which confers blasticidin resistance (Ozawa, 2018). After selection with blasticidin (8 μg/ml), colonies were isolated and tested for the expression of the gene products by immunofluorescence staining and Western blotting.

### Transfection of cells with siRNA oligos and expression vectors

Protein depletion was achieved using Stealth siRNAs (Invitrogen). The following oligos were used: NCKAP1 HSS116652 (#1), NCKAP1 HSS116650 (#2) and NCKAP1 HSS173885(#3) for Nap1; ARPC2 HSS173400 (#1), ARPC2 HSS115366 (#2) and ARPC2 HSS115367 (#3) for p34-Arc/ARPC2. We also used negative control siRNAs (ThermoFisher cat#12935-300; Invitrogen cat#46-2000). All siRNAs were transfected using Lipofectamine RNAiMAX Reagent (Invitrogen Cat#13778-150), according to the manufacturer’s protocol. The efficiency of protein depletion was verified by Western blots and immunofluorescence staining. Data obtained using one of the multiple siRNAs were shown as representatives, after confirming that others showed similar effects.

To construct Lifeact-RFP, we replaced the region between Sal1 and HindIII sites of the pCAH-LifeAct-GFP plasmid, which was published previously (Nishimura et al., 2016; Riedl et al., 2008), with the same region of pCANw-RFP plasmid. GFP-AHPH-DM (Tse et al., 2012) (Addgene plasmid #71368), as well as its control mutant version GFPAHPHA740D/ E758K-DM, were a gift from A. Yap (University of Queensland). Nap1-GFP was described previously (Hayashi et al., 2014). Wild-type and mutated MLC2, AA- and DDMLC2, tagged with EGFP at the C terminus were a generous gift of Shigenobu Yonemura (Tokushima University) (Watanabe et al., 2007). Plasmids were introduced into cells using Lipofectamine LTX Plus Reagent (Invitrogen Cat#15338-100), according to the manufacturer’s protocol.

### Antibodies and other reagents

We used the following primary antibodies: Mouse anti-β-catenin (BD Transduction Cat#610153, 1:100 for IF); rabbit anti-α-catenin (Cell Signaling Cat#8480S, 1:100 for IF); rabbit anti-a-catenin (Sigma Cat#C2081, 1:100 for IF); mouse anti-a-catenin (ENZO Life Sciences Cat#ALX-804-101-C100, 1:100 for IF); mouse anti-P-cadherin (TAKARA Cat#M127, 1:100 for IF); mouse anti-Rac1 (Abcam Cat#ab33186, 1:100 for IF); mouse anti-claudin 7 (Invitrogen Cat#37-4800, 1:100 for IF); rabbit anti-ARPC2 (Abcam Cat#ab133315, 1: 300 for Western blot); rabbit anti-p34-Arc/ARPC2 (EMD Millipore Cat#07-227, 1:100 for IF); mouse anti-p120-catenin (BD transduction Cat#610134, 1:100 for IF); mouse anti-ABI-1 (MBL Cat#D147-3, 1:100 for IF); rabbit anti-phospho-Myosin Light Chain 2 (Thr18/Ser19) (Cell Signaling Cat#3674S, 1:100 for IF); rabbit anti-Myosin IIA (Sigma-Aldrich Cat#M8064, 1:100 for IF); rabbit anti-Myosin IIB (Sigma-Aldrich Cat#M7939, 1:100 for IF); rabbit anti-Cortactin (Cell Signaling Cat#3503S, 1:100 for IF); rabbit anti-GFP (MBL Cat#598, 1:500-1000 for IF); rat anti-E-cadherin (ECCD2, ascites fluid, 1:100 for IF); mouse anti-E-cadherin (HECD1, ascites fluid, 1:100 for IF); mouse anti-a-tubulin (Sigma Cat#T9026, 1:500 for WB); and rabbit anti-phospho-Histone H3 (Ser10)(Sigma-Aldrich Cat#06-570, 1:100 for IF). Secondary antibodies use were: Alexa Fluor 488 donkey antimouse IgG (Life Technologies Cat#A21202); Alexa Fluor 594 goat anti-mouse IgG (Life Technologies Cat#A11032); Alexa Fluor 647 goat anti-mouse IgG (Invitrogen Molecular probes Cat#A21236); Alexa Fluor 488 goat anti-rabbit IgG (Invitrogen Cat#A11034); Alexa Fluor 594 goat anti-rabbit IgG (Life Technologies Cat#A11037); Alexa Fluor 647 goat antirabbit IgG (Invitrogen Molecular probes Cat#A21245); Alexa Fluor 488 chicken anti-rat IgG (Invitrogen Molecular probes Cat#A21470); and Alexa Fluor 594 donkey anti-rat IgG (Invitrogen Molecular probes Cat#A21209).

For actin staining, we used phalloidin-488 (Invitrogen Cat#A12379, 1:500), phalloidin-(Invitrogen Cat#A12381, 1:1000) and phalloidin-647 (Invitrogen Cat#A22287, 1:500). The following inhibitors were used: Blebbistatin (Sigma Cat#B0560); EHT1864 (Abcam Cat#AB229172); CK666 (MERK Cat#182515); and Latrunculin A (Sigma Cat#L5163).

### Immunofluorescence staining and microscopy

Cells were fixed with 1 to 2% paraformaldehyde (PFA) in PBS (pH 7.4) for 10 to 15 min, permeabilized with 0.25% Triton X-100 in PBS for 10 min, blocked for 30 min with 3% BSA in PBS, then incubated with primary antibodies (2 h), followed by incubation with secondary antibodies and/or phalloidin (1 h) in a blocking buffer. Three 10-min washes with PBS were performed after first and secondary antibodies incubations. After washing with distilled water, samples were mounted in FluorSave Reagent (CalbiochemCat#345789-20ML). All steps were carried out at room temperature. Samples were analyzed by Zeiss Axiopan 2 or Axio Imajor.Z2 through Plan-APOCHROMAT 63x/1.4 Oil Dic or Plan-NEOFLUAR 40x/1.3 Oil Dic objectives, or a laser scanning confocal microscope (LSM780) on an inverted Axio Observer.Z1 through Plan-A APOCHROMAT 63x1.4 Oil DIC objectives. Generally, we used conventional optical microscopes for photographing the marginal and sub-marginal zones of a colony, as they are thin, and confocal microscopes for photographing interior cells, which are thicker than marginal cells. Photographic images were processed with ImageJ/Fiji software.

### Time-lapse movies

For analysis of wound healing and cell migration by live imaging, we used a LCV100(Olympus) equipped with a UAPO x40/340 objective lens (Olympus), a LED light source, a DP30 camera (Olympus), differential interference contact (DIC) optical components and interference filters, except Video 2, for which we used an inverted fluorescence microscope (IX-81, Olympus, Japan) equipped with a spinning disk confocal imaging unit (CSU-X1, Yokogawa), a 40/1.35 oil immersion objective (UApo/340, Olympus) and EMCCD (iXon+, Andor Technology). To observe actin dynamics, we isolated stable lines of wild-type and αEcat KO Caco-2 cells expressing Lifeact-RFP. A wild-type line of these transfectants was additionally transfected with Nap1-GFP in a transient way. These cells were observed using an inverted fluorescence microscope (IX-81, Olympus, Japan) equipped with a spinning disk confocal imaging unit (CSU-X1, Yokogawa), a 40/1.35 oil immersion objective (UApo/340, Olympus), and a 561 nm laser (Sapphire LP, Coherent) for RFP excitation or a 488 nm laser (Sapphire LP, Coherent) for GFP excitation. We took fluorescence images with multiple zstacks by EMCCD (iXon+, Andor Technology) at the specified time intervals, and then made maximum intensity Z projections.

### Lattice light-sheet microscopy (LLSM)

For LLSM imaging, Caco2 cells expressing Lifeact-RFP were seeded on a collagen-coated coverslip 5 days before imaging. During imaging, cells were maintained in a L-15 medium (Sigma) supplemented with 10% serum at 25 °C. A LLSM was built in house following the design of the Betzig lab (Chen et al., 2014), with a slight modification. Among four cylindrical lenses we installed only f=25 nm and f=250 nm cylindrical lenses. For live imaging of Lifeact-RFP, a 560 nm laser (MPB Communications) and a long pass emission filter BLP02-561R-25 (Semrock) were used to acquire images through a CFI Apo LWD 25XW 1.1 NA detection objective (Nikon) and a sCMOS camera Orca Flash 4.0 v(Hamamatsu Photonics). To create a lattice light-sheet, a dithered square lattice was used through a spatial light modulator (Fourth Dimension Displays) in combination with an annular mask with 0.55 outer- and 0.44 inner-numerical apertures (Photo-Sciences Inc.), and a custom NA 0.65 excitation objective (Special Optics). Image stacks were collected with a 200 nm step size between planes with 20-msec per plane exposure times and 24.099-sec intervals. Post-acquisition, images were deskewed and deconvolved using LLSpy (Lambert and Shao, 2019). After deskew processing, the voxel pitch was 0.104×0.104×0.103 μm. Images were represented in 3D using Imaris software (Bitplane).

### Image analysis and quantification

Tracking of individual cells was performed with an Image/Fiji plugin, Manual Tracking. Intensity of immunofluorescence antibody stains was measured using MetaMorph Image Analysis Software, and the overlapping of the stains derived from different antigens were analyzed by a MetaMorph application, Measure Colocalization. The areas in an image were measured using Image/Fiji.

## Acknowledgements

We thank S. Ito for advice on time-lapse imaging, T. Kimura for help in statistical analysis, and the RIKEN Kobe light microscopy facility for imaging experiments. The lattice lightsheet microscope was home built in the Kiyosue lab at RIKEN Center for Biosystems Dynamics Research (BDR) under a research license agreement from Howard Hughes Medical Institute (HHMI), and we thank E. Betzig (HHMI, Janelia Research Campus) and W. Legant (HHMI, Janelia Research Campus) for their generous support by providing technical information and operational know-how. We also thank M. Yamaguchi (Carl Zeiss Microscopy Co., Ltd.) for support using the Imaris software. This work was supported by intramural funds of RIKEN BDR to M.T.; Grant-in-Aid for JSPS Fellows (Grant No. 18J01239) to T.Y.; and the Japan Society for the Promotion of Science–NEXT program (No. LS128), Takeda Science Foundation, the Uehara Memorial Foundation, a Grant-in-Aid for Challenging Exploratory Research (KAKENHI No. 18H05371), and JST CREST (No. 18H05371) to Y. M.-K. The authors declare no competing financial interests.

## Author contributions

M. Ozawa and M. Takeichi conceived and designed the study. M. Ozawa and S. Hiver performed most parts of the biological experiments and data analysis of wound healing experiments. T. Yamanoto performed fluorescence microscopy imaging presented in Videos 2, 3, and 5 to 11. T. Yamanoto and T. Shibata performed data analysis in the experiments presented in Fig. 1. Y. Mimori-Kiyosue performed lattice light-sheet microscopy and image processing. S. Upadhyayula provided critical advice on operation of LLSM. M. Takeichi analyzed the data and wrote the manuscript.

## Supplemental Figure Legends

**Figure S1.**
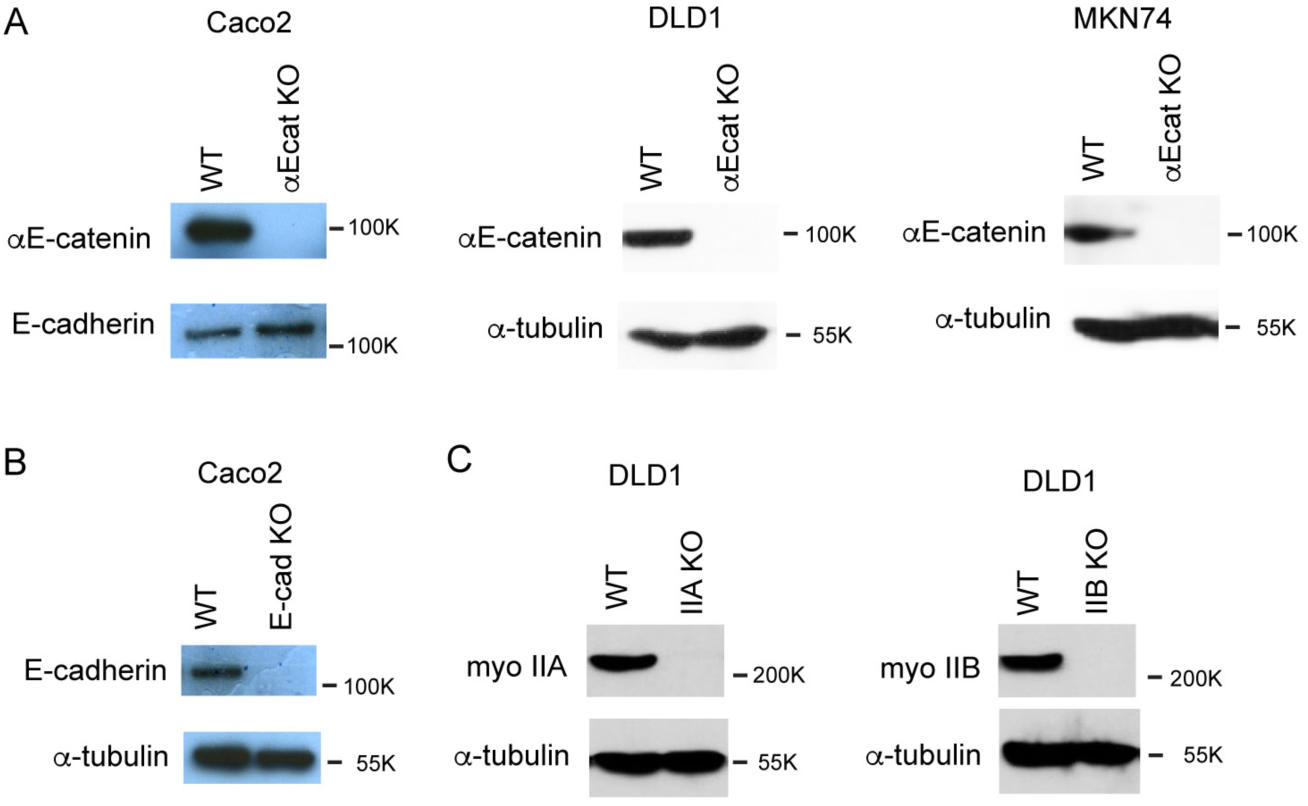
Western blotting analysis of CRISPR/Cas9-mediated deletion of proteins. (**A**) αE-catenin. (**B**) E-cadherin. (**C**) myosin IIA (myo IIA) and IIB (myo IIB).

**Figure S2.**
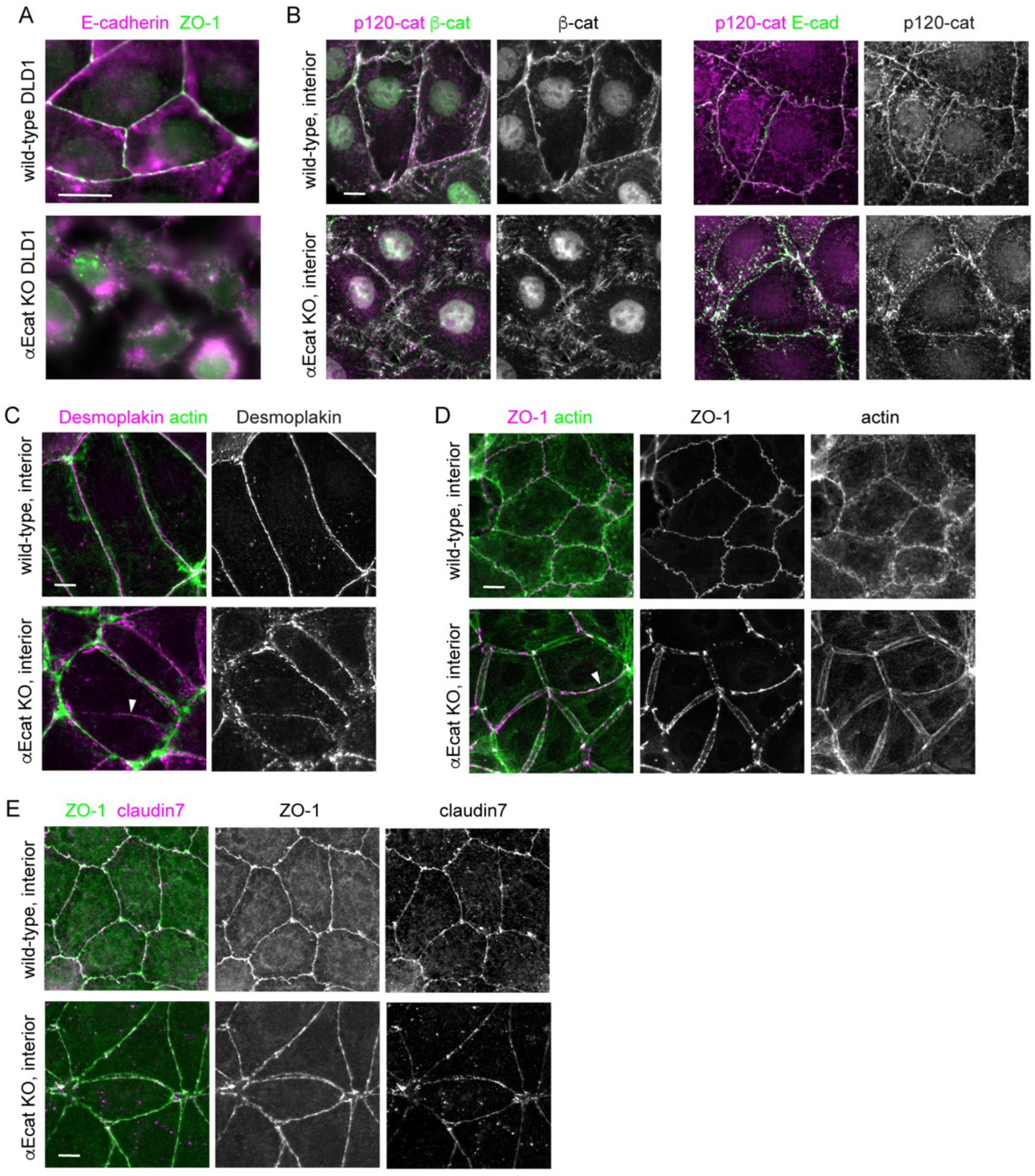
Junctional proteins in DLD1 and Caco2 cells. (**A**) Co-immunostaining for Ecadherin and ZO-1 in wild-type or αE-cat KO DLD1 cells. (**B** to **E**) Co-immunostaining for the indicated proteins in wild-type or αEcat KO Caco2 cells. Arrowheads indicate closed junctions in αEcat KO cells. p120, p120-catenin; β-cat, β-catenin. Scale bars, 10 μm.

**Figure S3.**
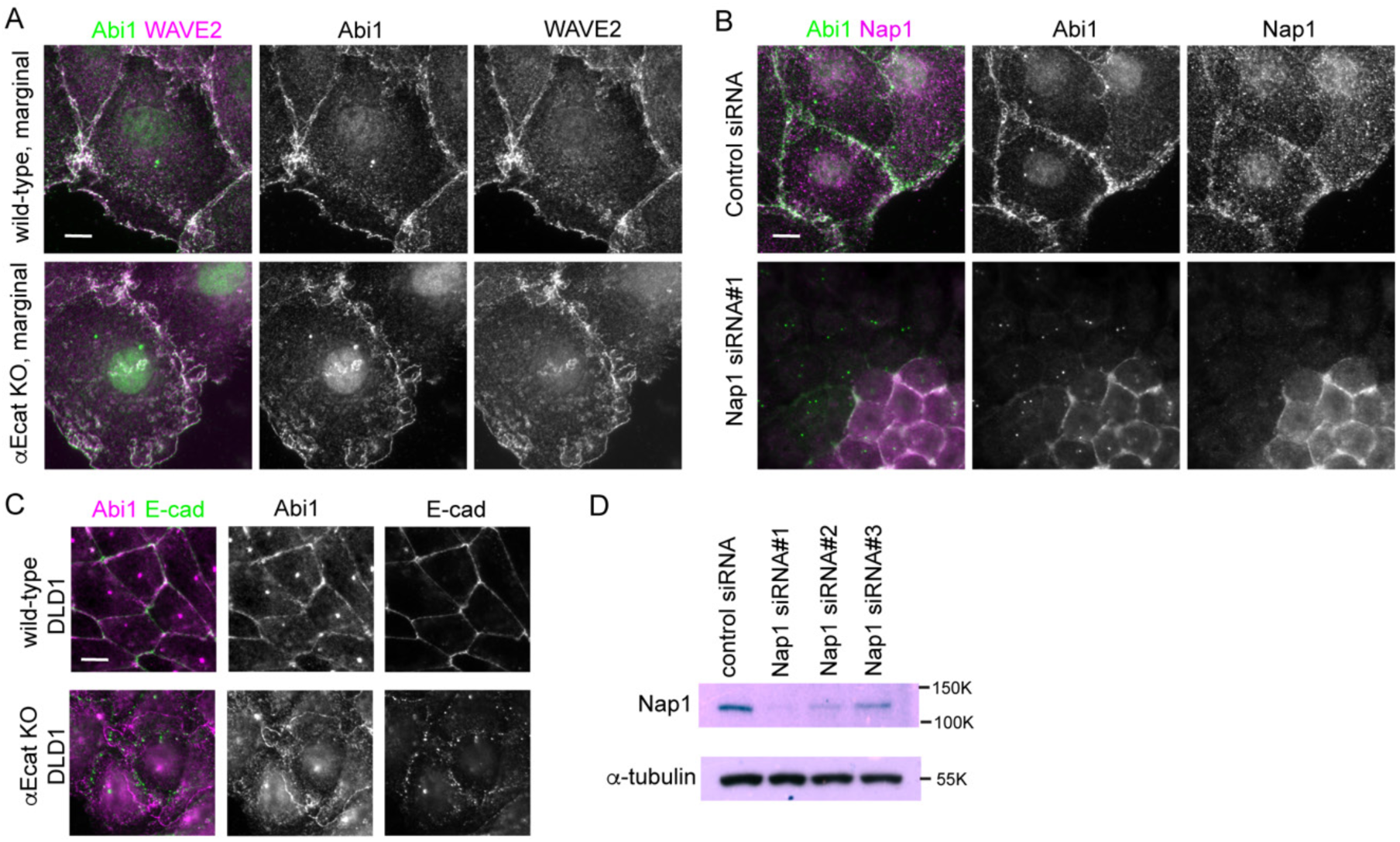
Detection of WRC components in Caco2 and DLD1 cells. (**A**) Coimmunostaining for Abi1 and WAVE2 in wild-type or αEcat KO Caco2 cells. (**B**) Coimmunostaining for Abi1 and Nap1 in wild-type Caco2 cells, which have been treated with siRNA for Nap1. Junctional Abi1 was removed as a result of Nap1 depletion, whereas centrosomal staining with anti-Abi1 antibody was not, which suggests that this staining is due to non-specific binding of the antibody to centrosomes. Consistently, antibodies for WAVEor Nap2 did not detect centrosomes. (**C**) Co-immunostaining for Abi1 and E-cadherin in wild-type and αEcat KO DLD1 cells. (**D**) Western blots for Nap1 in Nap1 siRNA-treated cells. Scale bars, 10 μm.

**Figure S4.**
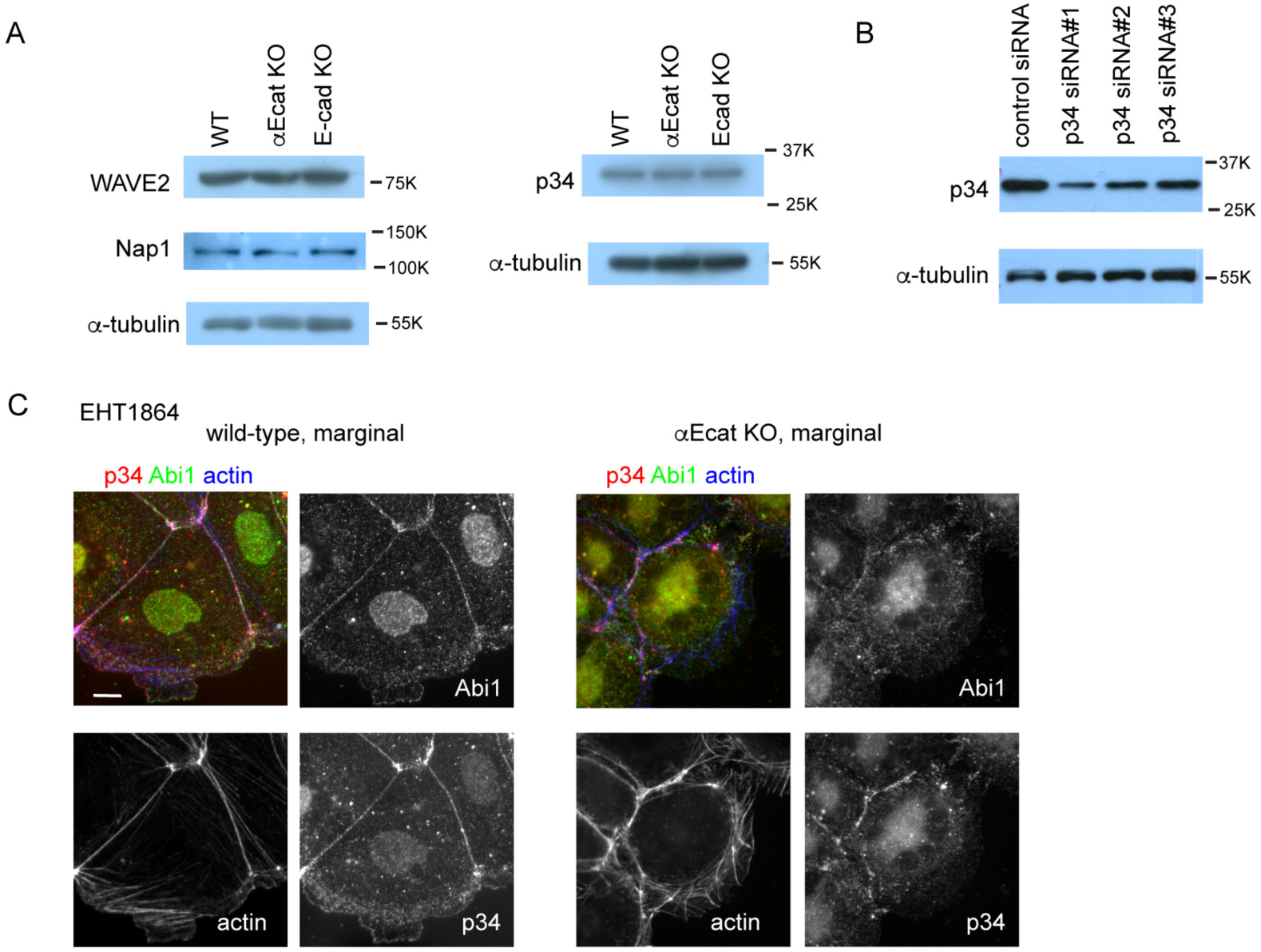
Expression of WRC and Arp2/3 complex, and effect of Rac1 inhibitor. (**A**) Western blots for WAVE2, Nap1 or p34/ARPC2 (p34) in wild-type, αEcat KO and E-cad KO Caco2 cells. The Nap1 blot was obtained separately from the WAVE2 and a-tubulin blots. (**B**) Western blots for p34/ARPC2 in p34/ARPC2 siRNA-treated cells. (**C**) Effect of the Rac1 inhibitor EHT1864 on actin and Abi1 distribution in wild-type or αE-cat KO Caco2 cells. Cells were incubated with 100 μM EHT1864 for 5 hr 30 min. Scale bars, 10 μm.

**Figure S5.**
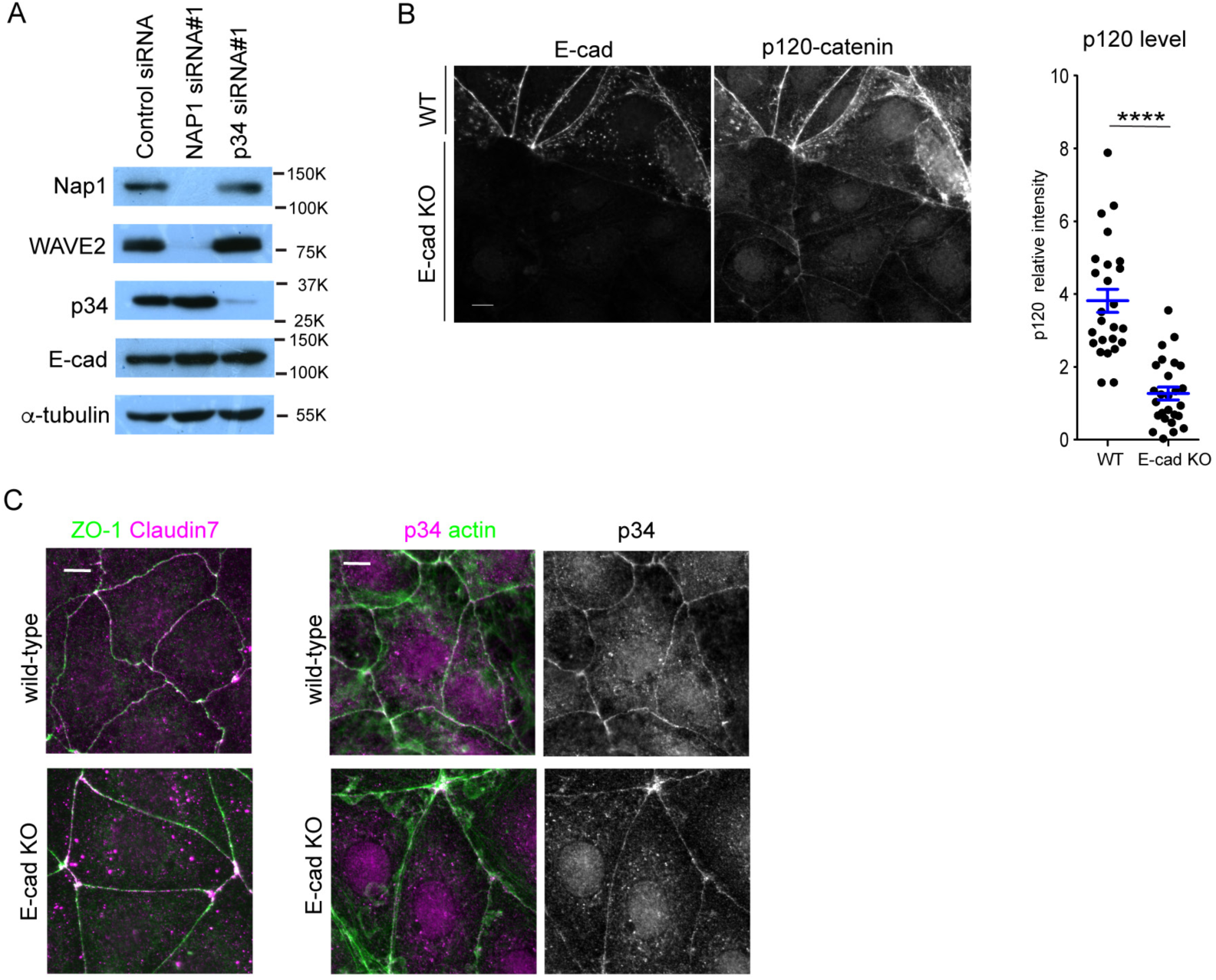
Effect of NAP1 or p34/ARPC2 depletion and E-cadherin deletion on other proteins. (**A**) Effect of Nap1 or p34/ARPC2 depletion on the level of other proteins. (**B**) Effect of E-cadherin deletion on p120-catenin level. Wild-type and E-cad KO Caco2 cells were mixed, and co-immunostained for E-cadherin (E-cad) and p120-catenin, which allows direct comparison of the level of these proteins between the two cell populations. Graph, relative immunostaining intensity of p120-catenin. A few points in each of the bi-cellular junctions were randomly selected for measurement, using 6 wild-type and 7 E-cad KO cells. ****p<0.0001 (t-test). (**C**) Co-immunostaining for ZO-1 and claudin 7, or 34/ARPC2 and actin in wild-type and E-cad KO Caco2 cells. Scale bars, 10 μm.

## Video Legends

**Video 1.** Time-lapse images of wild-type (left) and αEcat KO (right) DLD1 cells at the edge of a wound. The images were acquired at 10-min intervals, displayed at 12 fps.

**Video 2.** Time-lapse images of an isolated wild-type DLD1 cell. The images were acquired a 5-min intervals, displayed at 7 fps.

**Video 3.** Time-lapse images of a sheet of wild-type Caco2 cells, expressing Lifeact-RFP, at the wound edge. The images were acquired at 5-min intervals, displayed at 12 fps.

**Video 4.** 3D time-lapse images of wild-type Caco2 cells, expressing Lifeact-RFP, at a marginal zone of a cell sheet, collected with a LLSM. The video is replayed while being tilted. Time scale, h:m:s:ms.

**Video 5.** Time-lapse images of an isolated wild-type Caco2 cells, expressing Lifeact-RFP. The cell undergoes mitosis during imaging. The images were acquired at 5-min intervals, displayed at 12 fps.

**Video 6.** Time-lapse images of a small colony of wild-type Caco2 cells, expressing Lifeact-RFP. The images were acquired at 5-min intervals, displayed at 12 fps.

**Video 7.** Time-lapse images of a sheet of αEcat KO Caco2 cells, expressing Lifeact-RFP, at the wound edge. The images were acquired at 5-min intervals, displayed at 12 fps.

**Video 8.** Time-lapse images of an isolated αEcat KO Caco2 cells, expressing Lifeact-RFP. The cell undergoes mitosis during imaging. The images were acquired at 5-min intervals, displayed at 12 fps.

**Video 9.** Time-lapse images of a small colony of αEcat KO Caco2 cells, expressing Lifeact-RFP. The images were acquired at 5-min intervals, displayed at 12 fps.

**Video 10.** Time-lapse images of Nap1-GFP (green) and Lifeact-RFP (magenta) at a junction of wild-type Caco2 cells. The images were acquired at 5-min intervals, displayed at 6 fps.

**Video 11.** Time-lapse images of wild-type Caco2 cells not treated (left) or treated with μM CK666 (right). Imaging started at 14.5 hr after addition of the reagent. The images were acquired at 15-min intervals, displayed at 6 fps.

